# Chloroplast Methyltransferase Homolog RMT2 is Involved in Photosystem I Biogenesis

**DOI:** 10.1101/2023.12.21.572672

**Authors:** Rick G. Kim, Weichao Huang, Justin Findinier, Freddy Bunbury, Petra Redekop, Ruben Shrestha, TaraBryn S Grismer, Josep Vilarrasa-Blasi, Robert E. Jinkerson, Neda Fakhimi, Friedrich Fauser, Martin C. Jonikas, Masayuki Onishi, Shou-Ling Xu, Arthur R. Grossman

## Abstract

Oxygen (O_2_), a dominant element in the atmosphere and essential for most life on Earth, is produced by the photosynthetic oxidation of water. However, metabolic activity can cause accumulation of reactive O_2_ species (ROS) and severe cell damage. To identify and characterize mechanisms enabling cells to cope with ROS, we performed a high-throughput O_2_ sensitivity screen on a genome-wide insertional mutant library of the unicellular alga *Chlamydomonas reinhardtii*. This screen led to identification of a gene encoding a protein designated Rubisco methyltransferase 2 (RMT2). Although homologous to methyltransferases, RMT2 has not been experimentally demonstrated to have methyltransferase activity. Furthermore, the *rmt2* mutant was not compromised for Rubisco (first enzyme of Calvin-Benson Cycle) levels but did exhibit a marked decrease in accumulation/activity of photosystem I (PSI), which causes light sensitivity, with much less of an impact on other photosynthetic complexes. This mutant also shows increased accumulation of Ycf3 and Ycf4, proteins critical for PSI assembly. Rescue of the mutant phenotype with a wild-type (WT) copy of RMT2 fused to the mNeonGreen fluorophore indicates that the protein localizes to the chloroplast and appears to be enriched in/around the pyrenoid, an intrachloroplast compartment present in many algae that is packed with Rubisco and potentially hypoxic. These results indicate that RMT2 serves an important role in PSI biogenesis which, although still speculative, may be enriched around or within the pyrenoid.

**Significance Statement:** A high-throughput genetic screen was used to identify O_2_ sensitive mutants of *Chlamydomonas reinhardtii* (Chlamydomonas throughout) that experience elevated oxidative stress in the light relative to WT cells. Identification of genes altered in these mutants offers opportunities to discover activities that **a**) protect photosynthetic cells from oxidative damage, **b**) participate in rapid assembly of photosynthetic complexes, which would limit accessibility of intermediates to O_2_, and/or **c**) facilitate repair of damaged cellular complexes. A mutant from this screen disrupted for *RMT2*, originally described as encoding a Rubisco methyltransferase, was defective for PSI biogenesis. Additionally, RMT2 appears to be enriched in/around the pyrenoid, a chloroplast localized compartment harboring much of the Chlamydomonas Rubisco, raising the possibility that this compartment plays a role in PSI biogenesis.

## Introduction

In oxygenic photosynthesis, electron transport is driven by two photosystems, photosystem I and photosystem II (PSI and PSII). PSI uses absorbed light energy to drive electron transfer from plastocyanin (or cytochrome *c6*) to ferredoxin. Although the structure of PSI has been established at high resolution (1–6), there are still gaps in our knowledge of the steps involved in PSI biogenesis (7–11). Detailing PSI biogenesis has been challenging because PSI assembly is rapid, making it difficult to identify assembly intermediates (7, 8, 12), and the core subunits of PSI (PsaA and PsaB) constitute approximately 80% of the total molecular mass of the holocomplex. Rapid PSI assembly is considered essential as intermediate assembly products may generate reactive species that chemically modify and damage intracellular structures and activities (12). In addition, the rapid assembly of PSI may limit damage to certain prosthetic groups that are integral to PSI function; e.g. the three 4Fe-4S clusters of PSI are highly sensitive to O_2_ (13–15). Although O_2_ sensitive constituents of PSI may be protected from damage within the fully assembled holocomplex, they would likely be susceptible to damage as a result of accumulation of reactive oxygen species (ROS) in the light, particularly during the process of assembling the complex. CGL71, one of the nuclear-encoded factors required for normal PSI assembly, has been previously implicated in protecting assembly processes from oxidative disruption (9).

To identify proteins involved in protecting cells from O_2_ reactivity or in making the cells O_2_ sensitive when they are absent/aberrant, we performed a screen to identify O_2_-sensitive mutants of the unicellular photosynthetic eukaryote, *Chlamydomonas reinhardtii* (Chlamydomonas throughout). We used the entire population of mutants in the CLiP (<u>C</u>hlamydomonas mutant <u>Li</u>brary <u>P</u>roject) mutant library (16, 17) to perform a large-scale screen for O_2_ sensitive strains. All of the mutants in this library have unique built-in DNA barcodes that enable the use of deep sequencing for high-throughput quantification of the growth of each mutant in the library under different environmental conditions (17); in this case, under different light and oxic conditions (**Fig. S1**). This screen identified many strains potentially hypersensitive to ambient O_2_ levels relative to the WT, parental cells (**Fig. 1** and **Table S1**).

**Figure 1.**
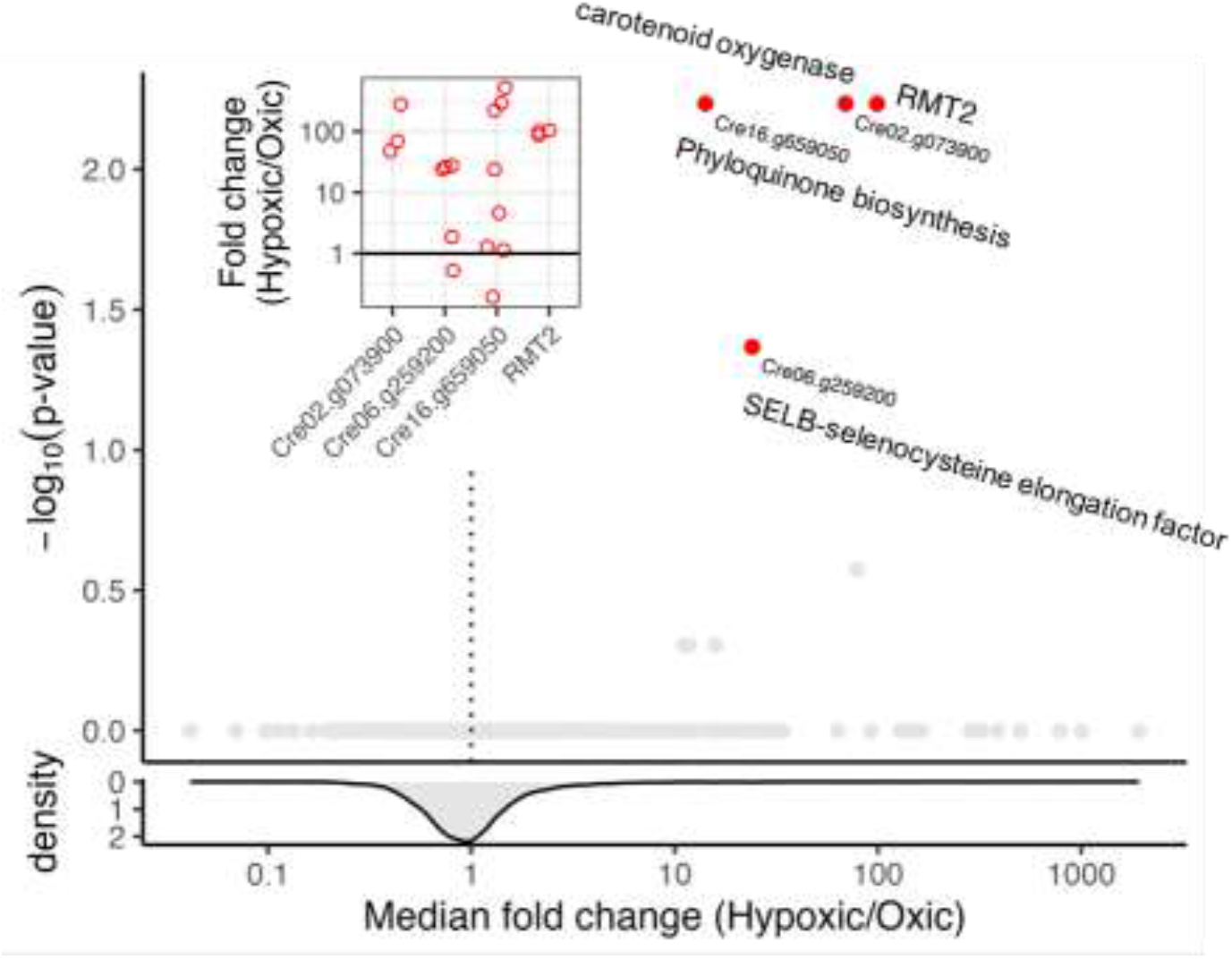
Several Chlamydomonas mutants survive better under hypoxic versus oxic conditions. Based on the barcode sequencing results, each barcode in the sample was assigned read counts. Several criteria were applied, sequentially, to filter the initial data counts as described in the **Materials and Methods**. The x-axis represents the median fold-difference in mutant counts+1 between hypoxic and oxic conditions for the mutant alleles. The density of the mutants, which mostly have a 1:1 ratio under HL hypoxic versus HL oxic conditions. The y-axis represents the BH-adjusted p-value. The points highlighted in red have an adjusted p-value<0.05 and are plotted in the top-left inset, which shows the Hypoxic/Oxic fold-change for each mutant allele. The protein names/functions are given in the figure and discussed in the text. The RMT2 identification is Cre12.g524500.

Two highly O_2_ sensitive mutants identified in the screen were disrupted for the gene encoding a homolog of Rubisco methyltransferase 2 (RMT2), which is thought to be involved in protein methylation, although this ‘RMT-like’ protein does not have all the typical features associated with methyltransferase activity. Post-translational modifications (PTM), including methylation of both histone and nonhistone proteins, play functional, regulatory, and structural roles in a range of organisms (18, 19). Advances in high-performance mass-spectrometry (MS) and quantitative proteomic approaches have enabled in-depth analyses of PTMs associated with proteins integral to photosynthetic function (20–23). The high sensitivity of *rmt2* mutants to O_2_ suggests that RMT2 activity and potentially protein methylation, can markedly impact processes that generate ROS, ameliorate damage by ROS, or participate in repair once damage occurs. Our additional analyses of the *rmt2* mutant suggest that it is aberrant for photosynthetic electron transport and is deficient in PSI, which generally leads to elevated levels of ROS in the light, causing phototoxicity (24, 25).

## Results

### Identification of *rmt2* mutant in O_2_ sensitivity screen

To screen for Chlamydomonas mutants sensitive to O_2_, we grew all mutants of the CLiP library as individual colonies on agar plates (grown in low light; LL), transferred them from the agar to liquid medium to generate a full library mutant pool, and then divided the pool for growth under oxic and hypoxic conditions in both LL and high light (HL) (**Fig. S1**). The O_2_ sensitivity of each mutant was individually assessed based on quantification of the number of barcoded reads representing each mutant in the library after pooling the cells and growing them in liquid medium under various growth conditions (17); each barcode is unique to a specific mutant. Growth differences for each mutant are presented as a fold change under hypoxic relative to oxic conditions following growth of the cultures for 7 generations at a light intensity of 300 µmol photons m^−2^ s^−1^. We show the hypoxic:oxic ratio and p-value for four of the mutant strains highly sensitive to O_2_ conditions in **Fig. 1**, with additional strains in **Table S1**. Strains mutated for the *RMT2* gene (Cre12.g524500) exhibited some of the highest O_2_ sensitivity scores among those mutants with high confidence differences (based on p-values) in their growth rates under hypoxic relative to oxic conditions (see **Materials and Methods**). Additional mutants represented in **Fig. 1** (see inset) and associated with O_2_ sensitivity identified in the screen encode carotenoid oxygenase (Cre02.g073900), which may be involved in the synthesis of strigolactone, a selenocysteine specific elongation factor/SELB (Cre06.g259200), and 2-succinyl-5-enolpyruvyl-6-hydroxy-3-cyclohexene-1-carboxylic acid synthase (Cre16.g659050), which is involved in the synthesis of the PSI cofactor vitamin K1 or phyloquinone. The strong spread of the hypoxic/oxic ratio for some mutants shown in the inset of **Fig. 1**, likely represents differences in the position of the inserted, disrupting cassette (e.g. in exon, intron, 5’ UTR). Other genes identified in the screen encode CGL71, a PSI assembly protein (9), RAA15 (Cre17.g728850) and RAT2 (Cre09.g388372), both components of the trans-splicing machinery involved in maturation of the *psaA* transcript (encodes core protein of PSI), and the GreenCut protein CTP2 (Cre10.g424775), a heavy metal transporting ATPase possibly required for chloroplast superoxide dismutase function (**Table S1**) (26).

In the two Chlamydomonas *rmt2* mutants, *rmt2-1* (CLiP catalog number: LMJ.RY0402.237082) and *rmt2-2* (LMJ.RY0402.255338), identified in this screen, the *APHVIII* gene (paromomycin resistance) was inserted into the 4th intron (*rmt2*-*1*) and 9th exon (*rmt2-2*) (**Fig. 2A**). These insertions result in a loss of gene function based on the phenotype of the mutants and the ability to rescue the mutant phenotypes by ectopic expression of the wild-type (WT) *RMT2* gene (see below). RMT2 has a predicted transit peptide at its N-terminal and both a Rubisco Substrate Binding (RSB) domain (**Fig. 2B**, underlined in blue) and a SET domain (acronym from the Drosophila Trithorax protein [Su(var)3-9, Enhancer-of-zeste and Trithorax]) (**Fig. 2B**, underlined in red); the latter domain harbors the active site of the enzyme that catalyzes the transfer of methyl groups from S-adenosyl-L-methionine (AdoMet) to the amino group of protein lysine residues, including those of histones (27). RMT2 also has homologs in *Pisum sativum*, *Glycine max*, *Spinacia oleracea*, and *Arabidopsis thaliana*, as shown in the Clustal Omega alignment of RMT2 homologs (**Fig. 2B**). Furthermore, Chlamydomonas has four other genes encoding proteins with significant homology to RMT2 (RMT1: Cre16.g661350; RMT3-5: Cre01.g017350, Cre08.g368700, Cre12.g503800, respectively). Examination of the sequences of these RMT homologs indicates that some of these proteins have the SET domain with an NHS motif, which is involved in AdoMet binding, as well as the specific catalytic tyrosine common to methyltransferase proteins of the RMT family (28). However, although RMT2 has a SET domain, this domain is missing the NHS motif and the catalytic tyrosine residue, although it does have a tyrosine that could potentially function in catalysis that is 9 amino acids upstream of the position of the catalytic tyrosine in other proteins of this family.

**Figure 2.**
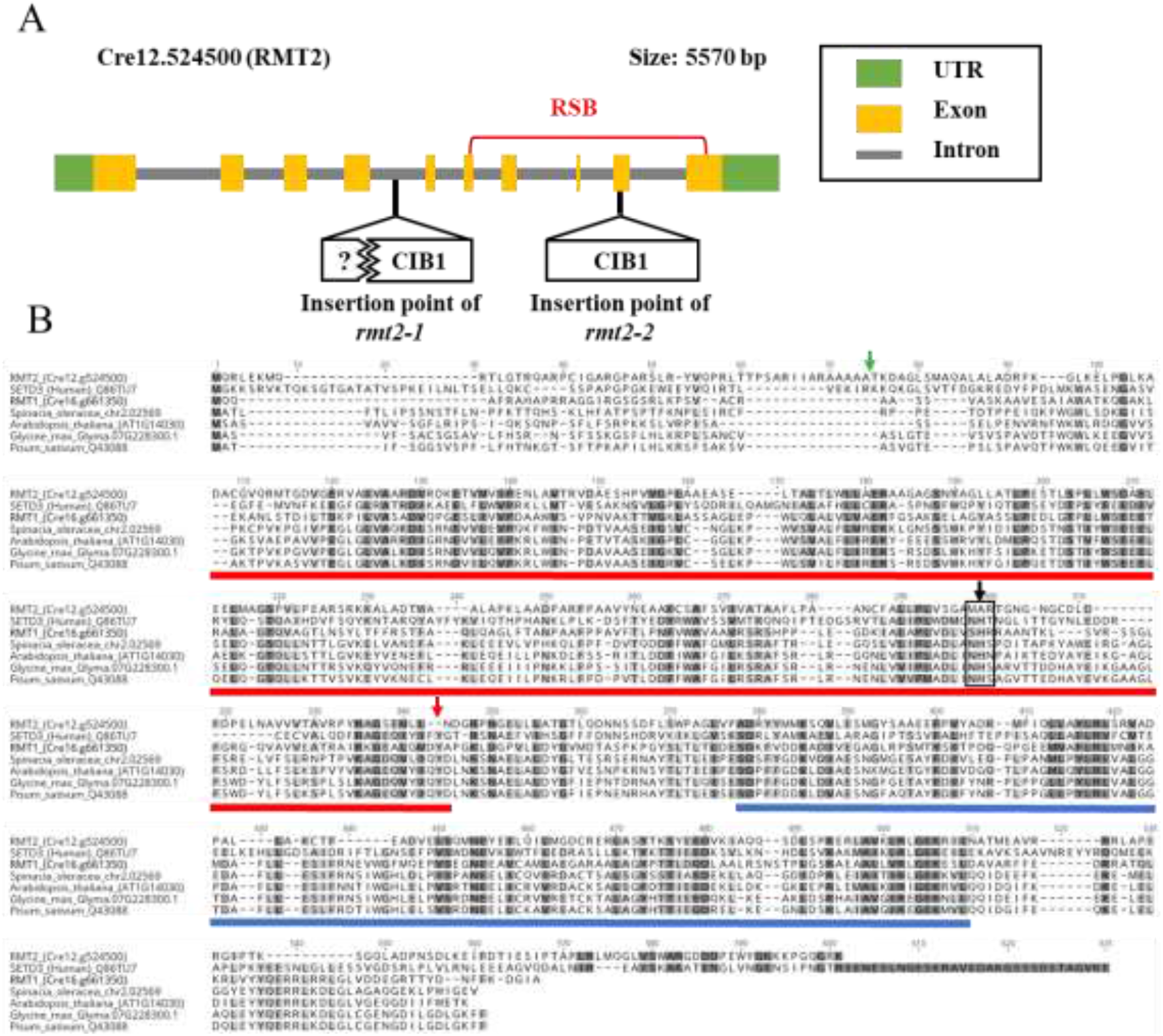
Positions of insertions in *rmt2* mutants, and RMT2 homologue sequence alignments. **(A)** *RMT2 g*ene showing position of inserted CIB1 cassette that includes *AphVIII* gene. Exons are *yellow* rectangles, introns *grey* lines, and 5’- and 3’-UTRs *green* rectangles. CIB1 is inserted into the 4th intron and 9th exon in the same and opposite orientation as the *RMT2* gene in *rmt2-1* and *rmt2-2* mutants, respectively. The junction sequences of the CIB1 insertion in *rmt2-2* were confirmed at both 5’ and 3’ ends. Only the 3’ junction was confirmed in *rmt2-1,* however, the sequence upstream of the 5’ end of *rmt2-1* 5’ is intact (from 11 bp upstream of the insertion site). The RSB (Rubisco Substrate Binding) domain of RMT2 is indicated in a red bracket. **(B)** Clustal Omega alignment of RMT2 homologs from *Spinacia oleracea, Arabidopsis thaliana, Glycine max,* and *Pisum sativum* along with SETD3 from humans and RMT1 from Chlamydomonas. The SET domain based on the SETD3 sequence is underlined in red. The RSB domain, based on the RMT2 sequences, is underlined in blue. The conserved residues are highlighted in dark grey (identical for all) and light grey (conserved for all). The NHS motif and the catalytic tyrosine residue of the SET domain in SETD3 are indicated by a black and red arrow, respectively.

### The *rmt2* mutant is high light sensitive and rescued by hypoxic conditions

Since *rmt2* mutants exhibited high O_2_ sensitivity based on our initial screen, they were expected to show improved survival under hypoxic conditions. Growth studies demonstrated that the *rmt2* mutants (here *rmt2-2*) were unable to grow under oxic, HL conditions (**Fig. 3A** and **FigS2**), but the growth rate of the mutant was similar to that of WT cells under hypoxic, HL conditions (**Fig. 3A**). Furthermore, growth of the *rmt2* mutants occurred under oxic conditions if the light intensity during growth was low; once the intensity was raised to above 50 µmol photons m^−2^ s^−1^ the cells stopped growing and lost viability (**Fig. S2A** and **S2B;** shown for *rmt2-1*). The O_2_ sensitive growth phenotype was rescued by ectopic expression of a WT *RMT2* gene in the *rmt2* mutants (**Fig. 3A**, *rmt2-2*(RMT2-NG)). These results demonstrate that both light and O_2_ levels strongly impact mutant growth and viability, raising the possibility that an aberration in photosynthetic processes in the mutant is responsible for the O_2_ sensitive phenotype.

**Figure 3.**
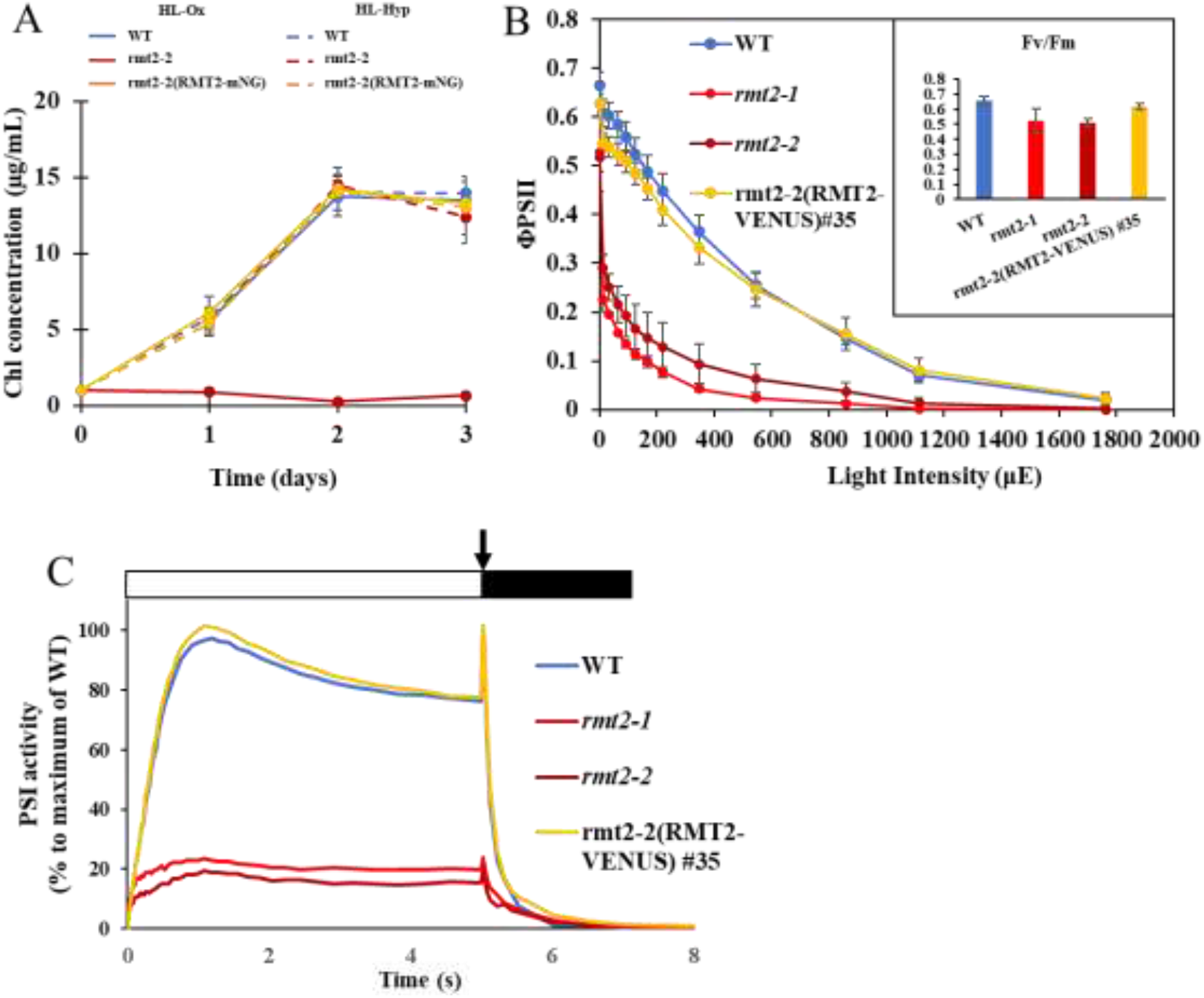
*rmt2* mutants are sensitive to oxygen and have aberrant photosynthesis. **(A)** Growth of WT and *rmt2-2* and rmt2-2(RMT2-mNG) under HL-Hy and HL-Ox conditions. WT, *rmt2-1* and rmt2-2(RMT2-mNG) cells were harvested during exponential growth in low light (30 µmol photons m^−2^ s^−1^) before inoculation into HL-Hy (750 µmol photons m^−2^ s^−1^; hypoxic gas*) and HL-Ox (750 µmol photons m^−2^ s^−1^; air) conditions. Cultures were bubbled with the indicated gas mixture at the flow rate of 500 mL min^−1^ with constant agitation (350 rpm). Chl concentrations were measured and plotted over the course of 3 days. *Hypoxic gas: 10% Air balanced by 89.96% N_2_ and 0.04% CO_2_. (µE = µmol photons m^−2^ s^−1^) **(B)** Quantum yield of PSII based on the fluorescence parameter ΦPSII, (which is (Fm′−Fs)/Fm′) in WT, *rmt2-1*, *rmt2-2*, and the *rmt2-2*(RMT2-VENUS) #35 rescued strains. Cells used for analyses were in exponential growth under low light (30 µmol photons m^−2^ s^− 1^) oxic conditions. For measurements, samples were incubated in the dark for 30 min and exposed to 1 min of actinic light at the intensities indicated on the x axis. Values are the mean of three biological replicates, and error bars represent standard deviations. The inset shows the Fv/Fm of each of the samples. **(C)** P700 oxidation and reduction kinetics of WT, the *rmt2-1* mutant, and the complemented strain (rescued with WT *RMT2* fused to a Venus tag). Cells were grown under LL-Ox conditions. Absorbance differences were monitored at 705 nm and 740 nm during continuous illumination with 300 µmol photons m^−2^ s^−1^ for 5 sec (white box), followed by a saturating light pulse at 1500 µmol photons m^−2^ s^−1^ (arrow) and a 3 sec dark incubation (black box). WT and *rmt2-1* samples were concentrated to 30 µg Chl mL^−1^. DCMU (PSII inhibitors) was included at a concentration of 10 μM to block electron flow from PSII. Kinetics were normalized by setting the maximum point of WT as 100%. Values are mean of three biological replicates.

### Photosynthetic activities are compromised in *rmt2*

To identify the cause(s) of light and O_2_ sensitivity, we characterized the photosynthetic activities of WT, the *rmt2-1* and *rmt2-2* mutants, and the *rmt2-2* rescued strains (using VENUS or mNeonGreen fluorophores). The allelic *rmt2* mutants both showed a marked reduction in ΦPSII relative to WT and the complemented cells at all light intensities, although the F_V_/F_M_ of the mutants relative to WT cells was only diminished by 10-20% (**Fig. 3B**, value after dark incubation, 0 µE). These findings suggest that PSII reaction center function in the mutant was not strongly compromised and that there is a lesion in photosynthetic function downstream of PSII. Measurements of PSI activity in *rmt2-1* and *rmt2-2* revealed a reduction by 80-90% of the activity measured in WT and complemented cells (**Fig. 3C**). This loss of PSI activity explains the diminished ΦPSII and HL sensitive phenotypes of the mutants. Immunoblot analysis of polypeptides of the photosynthetic apparatus, presented in **Fig. 4**, show that both mutants are strongly deficient in the levels of all PSI polypeptides examined (shown are PsaA, PsaC, PSAD, and PSAH; normalized to tubulin abundance) after growth under LL conditions (30 µmol photons m^−2^ s^−1^). Levels of most polypeptide subunits of other photosynthetic complexes, including ATP synthase (AtpA), cyt *b_6_f* (PetA), PSII (PsbA and PsbB), and Rubisco (RbcL and RBCS), were not strongly impacted in the mutants (**Fig. 4**), although some decline in the levels of the PsbA and PetA proteins was observed, suggesting that the function of RMT2 may extend beyond the accumulation of PSI, or that the mutant may experience some damage that impacts the levels of the other complexes of the photosynthetic apparatus even in LL. This potential secondary effect of lower PSI on the overall levels of other photosynthetic proteins is also consistent with the LC-MS/MS results in which PSI polypeptide levels are very low relative to WT cells (reduced by ∼80%) while proteins of the other complexes are less impacted (reduced by ∼30-40%) (**Fig. S3**). Interestingly, the chloroplast-encoded PSI assembly factors (Ycf3 and Ycf4) were significantly upregulated in both *rmt2* mutants (**Fig. 4**), raising the possibility that impaired PSI assembly in the mutants elicits a “partial” compensatory mechanism(s) that leads to elevated levels of PSI assembly factors, and/or that the cells accumulate more PSI assembly intermediates that maintain a sustained association with assembly factors, allowing for their accumulation.

**Figure 4.**
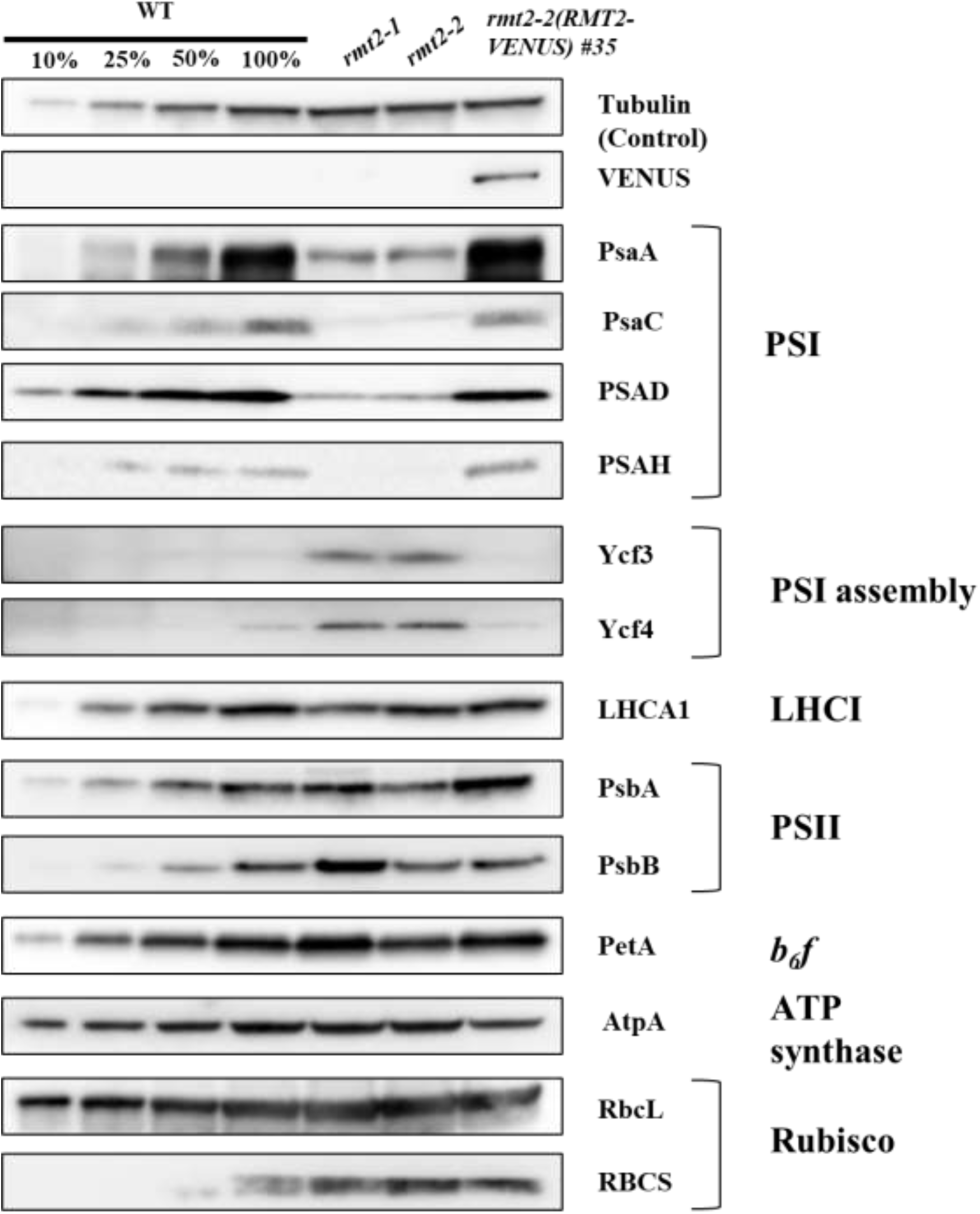
Immunoblot analysis of polypeptides of photosynthetic complexes of WT and *rmt2* mutants. Chlamydomonas proteins from WT, *rmt2-1* and *rmt2-2* grown under LL-Ox conditions were resolved by SDS-PAGE on a 12% polyacrylamide gel and detected immunologically. Antibodies used for the analysis were raised to polypeptides indicated on the right side of the figure. 1 μg of Chl was loaded for each sample analyzed by Western blots (except for the WT 10%, 25% and 50%, which were 0.1 μg, 0.25 μg and 0.5 μg, respectively).

### Impact of loss of RMT2 on protein profiles

To quantify the overall impact of a loss of RMT2 on the levels of Chlamydomonas proteins, we performed quantitative proteomics on WT and the *rmt2-1* and *rmt2-2* mutants after labeling cells in TAP medium containing ^15^N and ^14^N ammonium (reciprocal labeling experiments were performed using WT and the *rmt2-1* and *rmt2-2* mutant cells) under LL oxic conditions. Total proteins from ‘heavy’ and ‘light’ samples (see **Materials and Methods**) were extracted from the WT and mutant cells and equal amounts of the reciprocally labelled WT and mutant samples, based on chlorophyll (Chl) content, were combined, processed, and analyzed by mass spectrometry. A total of 4,573 proteins in all four of the samples were quantified (**Fig. 5A**). We identified 27 proteins, including HSP22F (Cre14.g617400), DEG11 Cre12.g498500), FTSZ1 (Cre02.g118600) and GPD2 (Cre01.g053000), that were elevated by at least 3-fold in the *rmt2* mutants relative to WT cells (**Fig. 5B, Table S2**). These proteins are associated with stress responses (protein folding and degradation) in Chlamydomonas, with GPD2 being involved in both acclimation to nutrient limitation and the synthesis of lipids (29). There were also 21 proteins that were at least 3-fold less abundant in the mutants relative to WT cells. Among them were many PSI-associated polypeptides (**Fig. 5B, Table S2**), as expected from our physiological and immunological analyses, and some CO_2_ concentrating mechanism (CCM)-related proteins, including CAH1, BST1 and others (**Table S2**). However, the absence of RMT2 in the cells does not alter the induction of CCM-associated genes under low CO_2_ conditions (**Fig. S4**) or the ability of the cells to maintain a prominent pyrenoid (**Fig. 6B**). The impact of RMT2 on the levels of CCM transcripts might be indirect and result from a higher intracellular accumulation of CO_2_ in the mutant during growth on acetate (acetate generates CO_2_ when metabolized) because of its reduced photosynthetic activity and inability to efficiently fix CO_2_ (resulting in elevated intracellular CO_2_ levels). This hypothesis is congruent with the findings that this mutant has a photosynthetic defect with reduced ΦPSII values, even at low light intensities (**Fig. 3B**), along with a marked reduction in PSI activity (**Fig. 3C**).

**Figure 5.**
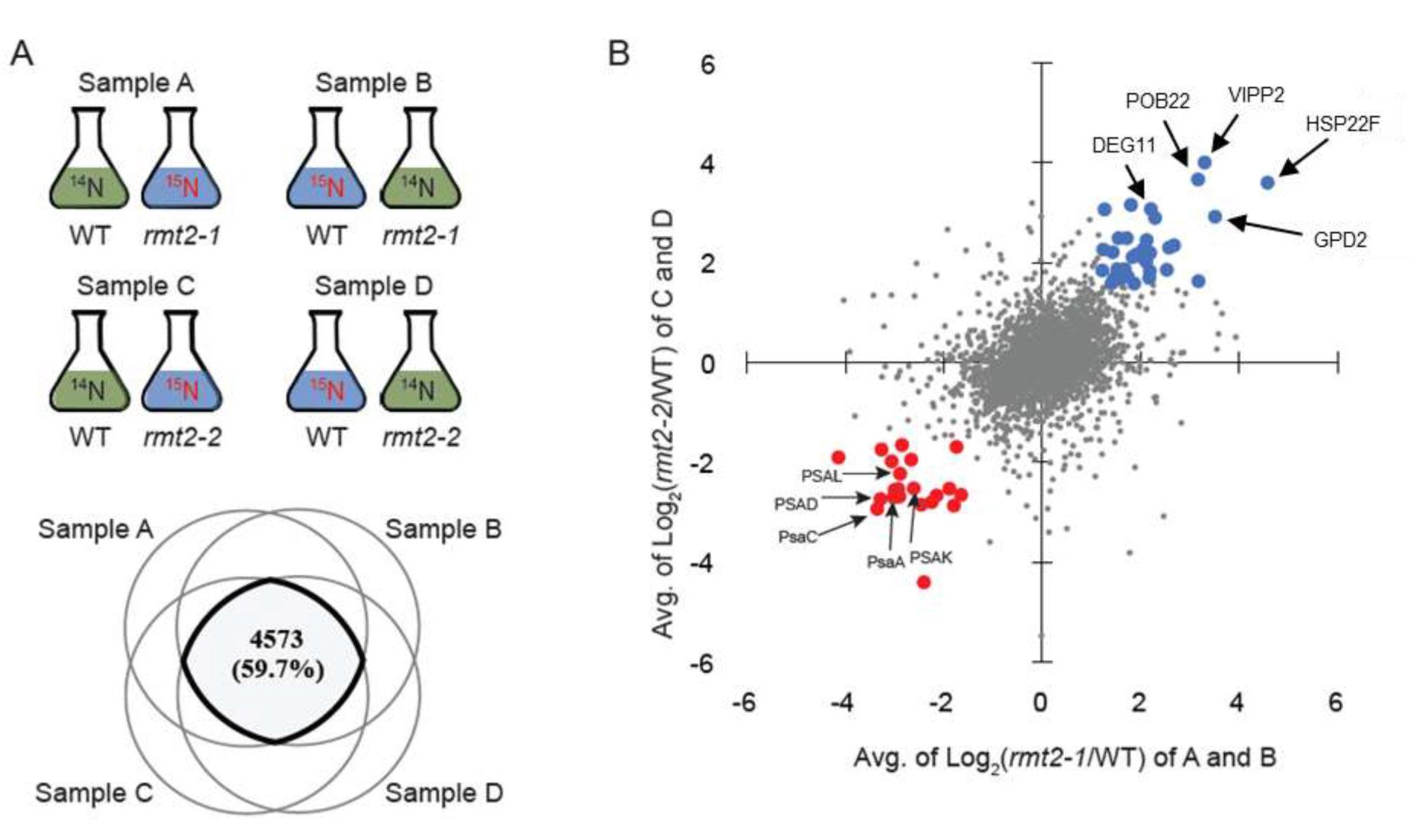
PSI family proteins show consistently lower protein levels in the *rmt2* mutant using quantitative analysis. **(A)** Schematic of 15N labeling and Venn diagram showing that 4573 proteins were detected and quantified in all four samples. **(B)** Plot showing the correlation between the *rmt2-1/*WT samples (sample A and B) and the *rmt2-2/*WT (sample C and D). Dots represent log2 values of the median ratios of *rmt2-1* vs WT of A and B on the x-axis and C and D on the y-axis. Those proteins that decreased by ≥3-fold in both *rmt2-1* and *rmt2-2* are highlighted in red, while those increased by ≥3-fold in both *rmt2-1* and *rmt2-2* are highlighted in blue. The remaining data points were marked as grey. PSI protein members (decreased level in *rmt2-1* relative to WT) and various stress related proteins (increased level in *rmt2-1* relative to WT) are labeled and their Cre numbers are given in **Table 2**.

**Figure 6.**
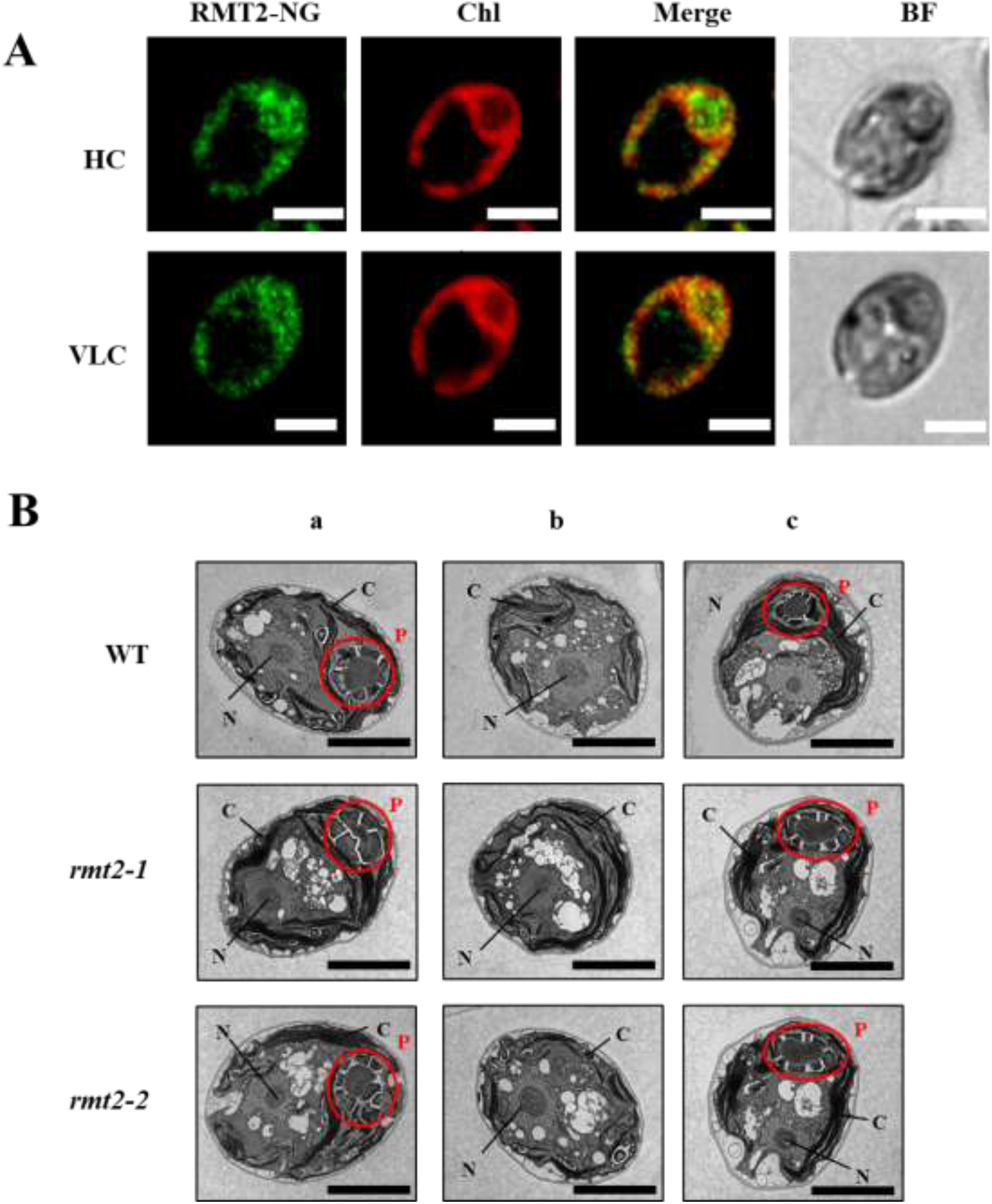
Subcellular localization of RMT2 and pyrenoid structure in *rmt2* mutants. **(A)** Localization of RMT2-mNG in high (HC, 3%) and very low (VLC, 4h at 0%) CO_2_ conditions. Shown are RMT2-mNG (mNeon green) localization, Chlorophyll autofluorescence (Chl), merge of Neon green and chlorophyll fluorescence (Merge), brightfield (BF). Each image is a maximal projection of 10 cross sections from the cortex to the middle of the cells. Scale bar: 5 µm. **(B)** TEM of sections of WT, *rmt2-1* and *rmt2-2* cells. (**a**) Cells with pyrenoid structures (grown in TAP medium at 20 µmol photons m^−2^ s^−1^). (**b**) Cells without pyrenoid structures (grown in TAP medium at 20 µmol photons m^−2^ s^−1^). (**c**) Cells with pyrenoid structures (grown in HS medium at 50 µmol photons m^−2^ s^−1^). Pyrenoid structures are highlighted by red circles. N, nucleus; C, chloroplast; P, pyrenoid. Scale bar = 5 μm.

### PSI defect in *rmt2* mutants appears to be at the post-transcriptional level

To test if the low PSI accumulation in the *rmt2-1* and *rmt2-2* mutants is a consequence of transcriptional regulation, we quantified transcript levels from both chloroplast and nuclear genes encoding subunits of PSI (*PsaA*, *PsaC*, *PSAD*, and *PSAE*) in WT, *rmt2-1, rmt2-2* and the rmt2-2(RMT2-Venus)#35 complemented cells grown under LL oxic conditions (**Fig. S5**). The chloroplast-encoded PSI genes (*PsaA* and *PsaC*) have similar transcript levels in the two *rmt2* mutants relative to the WT and the complemented cells whereas the nucleus-encoded PSI transcript levels (*PSAD* and *PSAE*) may be affected slightly by one of the mutant alleles (*rmt2-1*). The level of RMT2 transcripts detected in both *rmt2* mutants was very low. Overall, the results suggest that the lack of RMT2 does not strongly impact the level of PSI transcripts and that the function of RMT2 in PSI accumulation is likely post-transcriptional.

We also investigated if the reduced accumulation of PSI subunits in the *rmt2* background was due to their increased degradation rate (**Fig. S6**). In the presence of translation inhibitors of both chloroplast and cytoplasmic ribosomes (Chloramphenicol, CAP; Cycloheximide, CHX), the PSI subunit PsaA showed similar stability in the *rmt2-2* strain as in the WT background (compare top panels) and the complemented strain (bottom panel), suggesting that the diminished level of PSI in the RMT2 deficient strains is likely related to protein synthesis or less efficient assembly of PSI rather than the turnover rate of the assembled complex. PsbA protein turnover was used as a control, showing that the presence of CAP and CHX resulted in a more rapid loss of the PSII reaction center protein PsbA in WT cells, *rmt2-2* mutant cells and the rescued strain (*rmt2-2* + RMT2-mNG) relative to the untreated sample.

### RMT2 location under high and very low CO_2_ conditions

To help predict the function of RMT2, we determined its subcellular location under high and very low CO_2_. *RMT2* was fused to the sequence encoding the mNeonGreen fluorophore (RMT2-mNeonGreen) and the fusion construct was introduced into the *rmt2*-*2* mutant background (see **Materials and Methods**). The *rmt2*-*2* mutant phenotype was rescued by the RMT2-mNeonGreen construct; the transformed strain was able to grow under both HC (3%) and very low CO_2_ (VLC; <0.02% CO_2_) conditions (in TP medium at 150 µmol photons m^−2^ s^−1^). Under both conditions, much of the fluorescence of the RMT2-mNeonGreen fusion protein was associated with or in the vicinity of the pyrenoid, although there was a substantial amount that was more generally distributed in the chloroplast stroma (**Fig. 6A** and **Fig. S7**). Both the presence of a transit peptide and the association of RMT2 with proteins of the chloroplast (21) already suggested its localization in the chloroplast. However, these results also raise the possibility that the association of RMT2 with the pyrenoid may be important for the biogenesis of photosynthetic complexes, especially for PSI under conditions that would lead to the generation of strong oxidants in the stroma. Interestingly, the PSBP4 protein, which is involved in PSI assembly (30–32), also localized around the pyrenoid and is associated with pyrenoid tubules (33), and both RMT2 and CGL71 have been identified as interacting partners of PSBP4 (21). These results again support a role for RMT2 in PSI assembly and suggest that the process of assembly may occur in both the stroma and in association with the pyrenoid. Additionally, RMT2 localization did not appear to be impacted by the level of CO_2_ in the environment (**Fig. 6A**).

### Direct detection of RMT2 proximal proteins

To investigate which proteins are located proximal to RMT2 in vivo, we fused RMT2 with the engineered BioIDG3 (34) and transformed this construct, designated RMT2-ID, into the *rmt2-1* and *rmt2-2* mutants. The physiological function of the fusion proteins was confirmed by rescue of the mutant phenotype (growth in HL under oxic conditions). RMT2-VENUS expressed in the *rmt2-1* and *rmt2-2* mutants was used as a control. Using a similar ^15^N-labeling strategy (**Fig. 7A**), labeled cells were treated with biotin (4 h) and harvested. Proteins were extracted and subjected to trypsin digestion. The resulting peptides were enriched for those that were biotinylated, followed by detection and quantification of these peptides (**Fig.7B**) (35, 36).

**Figure 7.**
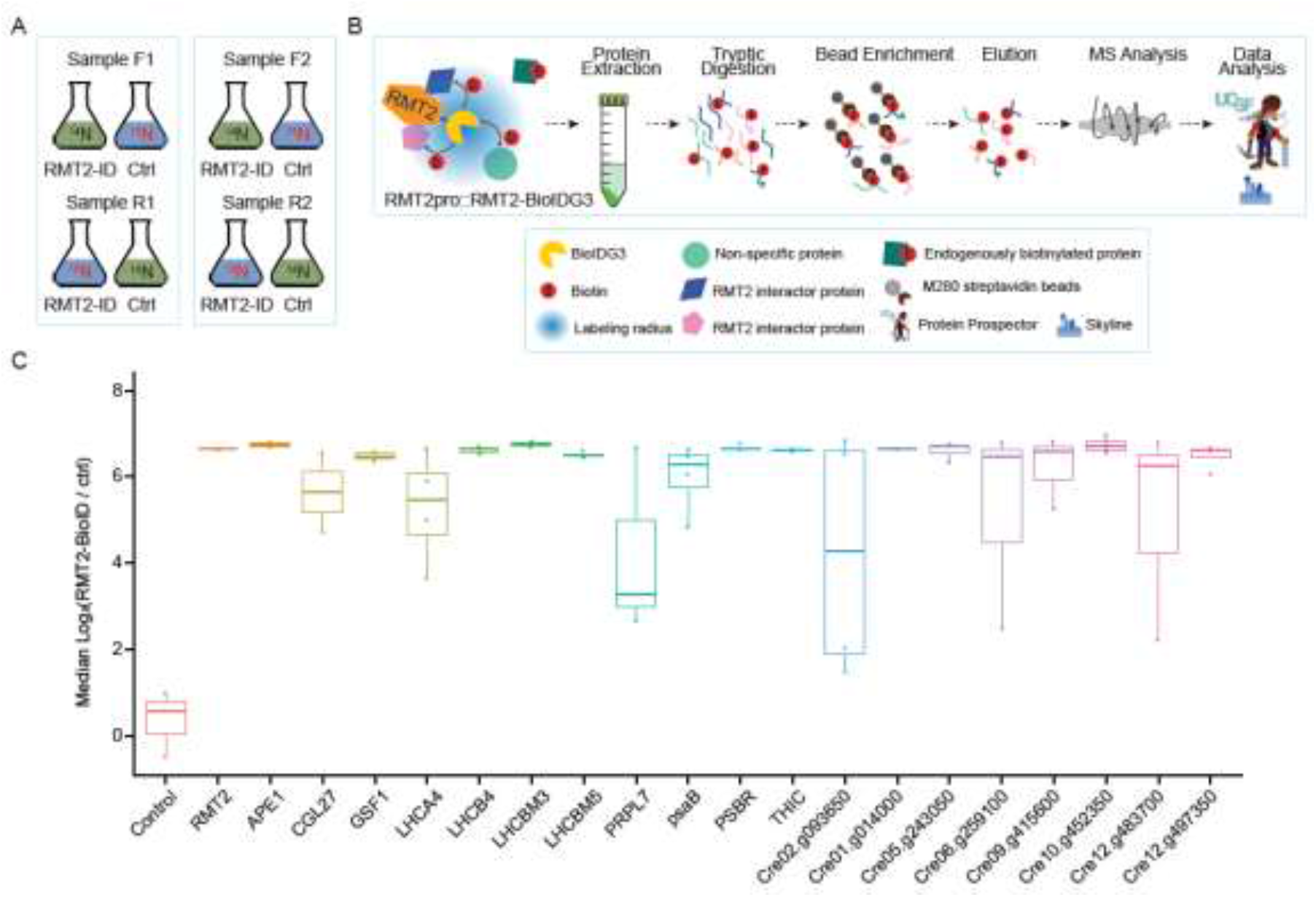
RMT2-ID enriches 19 specific proximal proteins, including PsaB and others. **(A)** SILIA experimental design included two RMT2-BioID forward- and reverse-labeled samples in *rmt2-1* background (set 1) and two RMT2-BIoID forward- and reverse-labeled samples in *rmt2-2* background (set 2). **(B)** Schematic workflow for enrichment of biotinylated peptides. Proteins are extracted and digested, and resulting peptides are subjected to streptavidin bead enrichment for biotinylated peptides and quantified using the ^15^N workflow. **(C)** RMT2 bait and a number of thylakoid-localized proteins are enriched by RMT2-BioIDG3, including psaB protein.

RMT2 was significantly enriched in the biotinylated fraction, with multiple biotinylated peptides from RMT2 identified (**Table S3**), indicating that the BioIDG3 was working in Chlamydomonas. In addition, 19 other biotinylated proteins are enriched, with a number of proteins predicted to be thylakoid membrane proteins, including several members of the chlorophyll A/B binding protein family (LHCA4, LHCB4, LHCBM3 and LHCBM5), the PSI subunit PsaB, and PSBR which is involved in the binding of LHCSR3 to the antenna complexes (**Fig. 7C**). These results suggest that RMT2 associates with thylakoid membranes and is proximal to many proteins involved in photosynthetic electron transport (associated with both PSI and PSII).

### Methylation of subunits of the photosynthetic apparatus

To investigate whether the PSI complex that accumulated in the *rmt2* mutants showed any aberrations in the methylation of PSI subunits, we purified thylakoid membranes from WT, *rmt2-1* and *rmt2-2* and quantified the methylated peptides of the different PSI subunits (PSAD and PSAE). While more than 50 methylated proteins of the thylakoid membranes were noted, those with very high confidence methylation are listed in **Table S3**. In addition, preliminary evidence suggests that there is no clear difference in the methylation of these thylakoid proteins in WT and the *rmt2* mutants, possibly indicating that the intermediates in PSI assembly with altered methylation are rapidly degraded or that methylation in the mutant is partially compensated for by other methyltransferases in the chloroplast.

## Discussion

We exploited the power of the CLiP library to simultaneously screen 58,101 mutants covering ∼83% of the Chlamydomonas nuclear protein-coding genes to identify strains more sensitive to oxidative conditions than the parental cells (16, 17). This analysis yielded a list of genes encoding proteins (**Fig. 1, Table S1**) that potentially allow Chlamydomonas to effectively cope with oxidative stress. The list includes CGL71, a protein previously shown to be required for protection of PSI assembly against oxidative damage (9) and proteins associated with the maturation of the mRNA encoding PsaA (RAT2 and RAA15) and PsaB; these findings establish the feasibility of this screen for obtaining mutants more susceptible to oxidative stress, such as mutants defective for PSI. Furthermore, this list provides an opportunity for researchers to discover novel proteins involved in photoprotection, and to identify O_2_ sensitive steps in PSI assembly as well as in repair of ROS damaged complexes.

RMT2 strongly impacts PSI accumulation (**Fig 3C** and **4**). The reduced level of PSI in the mutants results in O_2_ sensitivity (**Fig. 1** and **3**), similar to the phenotype observed for other PSI mutants (9, 37, 38). *rmt2* mutants exhibited reduced PSI activity under both LL oxic and dark oxic conditions (**Fig. S8**), which reflects a marked reduction in the levels of PSI subunits in these strains. These mutants die under HL oxic conditions but can be rescued (e.g. restores PSI activity, normal levels of PSI polypeptides and growth under HL oxic conditions) when a WT copy of the *RMT2* gene is ectopically expressed in the mutant genetic background (**Fig. 3C** and **4**). The mutant also grows well in HL under hypoxic conditions (**Fig. 3**), with PSI activity somewhat higher than under dark-oxic or LL-hypoxic conditions (**Fig. S8**); the activity can reach as much as ∼ 40% of that of WT activity under the former conditions. These phenotypes are different from those of the *cgl71* mutant, which accumulates near normal levels of PSI under dark-hypoxic conditions and almost no PSI when maintained under dark-oxic conditions (9). Hence, exposure of *cgl71* to ambient levels of O_2_ appears to inhibit PSI assembly in both the light and dark, which can be reversed when the cultures are made hypoxic. In contrast, in the *rmt2* mutant, PSI levels remain low in the dark, LL oxic and LL hypoxic conditions, with only a small increase in the level under HL hypoxic conditions, suggesting that the defect in PSI accumulation is not impacted strongly by oxic conditions, but that growth and viability are impacted. Additionally, hypoxia allows growth of *rmt2* under HL conditions without restoring WT levels of PSI. Therefore, RMT2 and CGL71 likely function at different stages of the assembly process, which in turn leads to different phenotypes; while the lower levels of PSI in *cgl71* can be mostly reversed under hypoxic conditions, in *rmt2* the PSI accumulation is markedly reduced even under hypoxic conditions, but the level of PSI maintained (15-20% under dark oxic and LL hypoxic and up to 40% under HL hypoxic conditions) allows the cells to grow as long as the O_2_ levels remain low. Furthermore, we were unable to show a difference in the rate of turnover PSI subunits in the *rmt2* mutant relative to WT cells, suggesting that the PSI complexes formed in the mutant cells are stable. In summary, whereas PSI biogenesis is affected by O_2_ in the *cgl71* mutant (tetratricopeptide repeat protein), possibly through protection of the PSI 4Fe-4S centers during assembly of the complex, as previously discussed (9), O_2_ does not strongly impact the biogenesis/accumulation of PSI in *rmt2*, but it does markedly influence its growth and viability in the light, suggesting that the growth defect is the result of the damaging impact of ROS generated by the photosystems, with ROS damage especially severe under HL conditions and when there is an imbalance in electron transport (39, 40).

Trimethylation of the ε-amino group of Lys-14 of the large subunit of Rubisco of tobacco and muskmelon was demonstrated over three decades ago (41). The enzyme associated with this modification was designated RMT (Rubisco methyltransferase), suggesting that the substrate of this enzyme may be RbcL. The function of RMT2 was predicted based on homology to Rubisco LSMT (<u>L</u>arge <u>S</u>ubunit <u>M</u>ethyltransferase) proteins from various plant species including *Pisum sativum* (pea), *Nicotiana rustica* (tobacco), *Spinacia oleracea* (spinach), and *Arabidopsis thaliana* (Arabidopsis). As shown in **Fig. 2**, RMT2 in Chlamydomonas is similar to plant RMTs and paralogous sequences in Chlamydomonas, such as RMT1, which encodes a methyltransferase that appears to be involved in methylation of fructose-bisphosphate aldolase (42). The RMT enzyme was initially purified and characterized from tobacco and pea (43–45). It is one of the few early non-histone lysine methyltransferases identified in plants. Most homologous proteins in other organisms, including those identified in non-plant species, were also given the name Rubisco LSMT, even though it was not established that the homologs functioned in methylation of RbcL. Interestingly, Lys-14 of the large subunit is conserved but not methylated in Chlamydomonas, spinach, or Arabidopsis (42, 46, 47). Therefore, RbcL is unlikely to be the substrate of LSMT proteins in Chlamydomonas, although RMT2 has an LSMT binding domain that may function in binding interacting partners. RMT type methyltransferases generally have SET domains that contain the catalytically critical residues identified in similar methyltransferases from other organisms (48). Specifically, the alignment in **Fig. 2B** shows that the highly conserved catalytic tyrosine residue and the NHS motif involved in binding S-adenosine methionine are present within the SET domain (28). These critical features are absent from Chlamydomonas RMT2, although there is a tyrosine residue in the Chlamydomonas protein that is 9 amino acids upstream of the tyrosine conserved in other organisms.

Since the level of transcripts from the PSI genes are not strongly impacted in *rmt2* (**Fig. S5**), and the mutant doesn’t appear to be affected in the turnover rate of PSI subunits (**Fig. S6**), the RMT2 protein is likely either directly or indirectly involved in translational or post-translational regulation of PSI biogenesis. Previous work showed that several photosynthetic processes are modulated by post-translational modifications. These processes include state transitions, the PSII repair cycle, control of translation through feedback control of CES proteins (49–51), and carbon assimilation (22). With respect to PSI biogenesis, little is known about the role of PTMs, although PSI subunits appear to be methylated based on our data (**Table S3**). In this study we found that protein subunits of other photosynthetic complexes (e.g. ATP synthase) of Chlamydomonas are also methylated (**Table S3**). Despite the widespread occurrence of methylation on chloroplast proteins of Chlamydomonas, there is little known about the specific enzymes involved in methylating these proteins or the functional consequence of this modification, which may cause conformation changes that alter biophysical and biochemical properties of the complexes (52). One of the previously characterized putative methyltransferases in Chlamydomonas is CIA6, which is important for pyrenoid formation; mutants of CIA6 exhibit altered pyrenoid morphologies under low CO_2_ (53). However, methyltransferase activity of CIA6 has not been experimentally confirmed, nor has its substrate been identified. Based on the use of fluorescently labeled fusion proteins, RMT2 localizes to the chloroplast and is potentially enriched in the pyrenoid. Importantly, *rmt2* mutants can still form pyrenoids (**Fig. 6B**) whereas *cia6* mutants cannot (53). These results suggest that while CIA6 and RMT2 potentially both have functions in the methylation of proteins, their downstream targets and molecular roles may be functionally distinct. Furthermore, we identified Cre08.g378250, which encodes another methyltransferase required for protection of cells under oxic conditions, although its subcellular location is not known. An additional methyltransferase, SMM7, was shown by others to be potentially pyrenoid localized (33).

WT RMT2 protein fused to mNeonGreen was able to rescue the PSI deficient phenotype of the *rmt2* mutant. The introduced fusion protein localized to the chloroplast with a strong signal associated with the pyrenoid (**Fig. 6**). The *rmt2* mutant also exhibited elevated accumulation of the PSI assembly factors Ycf3 and Ycf4 (**Fig. 4**) and both CGL71 and RMT2 appear to interact with PSBP4 (21), another PSI assembly factor associated with the pyrenoid (30). The pyrenoid is an organelle specifically found in algae and hornworts and is important for CCM function (54, 55). It is traversed by thylakoid membrane tubules that penetrate the Rubisco matrix and is surrounded by a peripheral starch sheath. These three distinct pyrenoid compartments may be physically associated through Rubisco binding motifs (56). Chloroplast stromal localized thylakoids perform photosynthetic electron transport and O_2_ evolution, which would result in an oxic stroma that can generate ROS (57). A study performed over 30 years ago suggested that thylakoid membranes that penetrate the pyrenoids in some algae (including Chlamydomonas) contain active PSI but no or little PSII (58, 59), with a more recent proteomic study showing that this plastid structure harbors several PSI subunits with only a single detected PSII subunit (60). Furthermore, recent spatial-interactome results revealed enrichment of PSI subunits in the pyrenoid while PSII subunits of the pyrenoid were only partially assembled or inactive (21, 55). Furthermore, the pyrenoid is separated from the rest of the stroma by a starch barrier that is likely to limit O_2_ permeability. ROS generated in HL or conditions of nutrient deprivation may trigger changes in gene expression that are responsible for altering the pyrenoid structure and further physically isolating it from O_2_. A peripheral pyrenoid mesh may enhance its physical compartmentalization and create the ‘cement’ that holds the starch plates in place, which may allow it to serve as a more effective diffusion barrier (61–63). Additionally, recent proteomic work found that proteins involved in chloroplast translation may have physical associations with the pyrenoid (60, 64).

Recently it was discovered that despite the presence of high external inorganic carbon (5 mM bicarbonate), treatment of Chlamydomonas with H_2_O_2_, which acts as a ROS signaling molecule (65), significantly increased the pyrenoid size, Rubisco relocation to the pyrenoid, and the formation of tight, thick starch plates held together by a peripheral mesh (61). Aspects of the biogenesis of photosynthetic complexes can be susceptible to ROS damage, especially if Fe-S complexes are integral to their function. Therefore, we speculate that the pyrenoid may be a hypoxic micro-compartment serving as a site for the biogenesis of photosynthetic complexes under conditions when stromal oxidative conditions would compromise the assembly process. Hence, the pyrenoid may not only provide protection from photorespiratory carbon metabolism catalyzed by Rubisco, but also may function to protect O_2_ sensitive processes such as PSI assembly (9, 11). These results are congruent with RMT2 being enriched in/around the pyrenoid where it may interact with PSI assembly factors such as PSBP4; the latter was shown to localize to thylakoid lumen puncta in the stroma and around the pyrenoids (21, 31). Additionally, mutants unable to assemble the pyrenoid (*pyr^−^*) have been identified (66, 67). Such mutants are defective for the CCM, but the impaired photosynthesis observed when the mutant cells are maintained at low levels of CO_2_ can be rescued upon exposing them to high CO_2_ concentrations; the mutants under these conditions show normal thylakoid morphology with little difference in various photosynthetic activities (66). However, WT Chlamydomonas cells do not completely eliminate the pyrenoid when the CCM is repressed, suggesting that it may play roles in processes other than the CCM (64). Thus, while the pyrenoid is not essential for PSI assembly (or assembly of other photosynthetic complexes) under high CO_2_ conditions, such a role may only be unmasked when the oxidative conditions of the chloroplast stroma impact processes susceptible to oxidative disruption (e.g. assembly processes); in most instances this would occur under conditions that would induce the CCM, for which the pyrenoid is absolutely required. Furthermore, analysis of potential RMT2 interacting proteins based on proximity labeling under (**Fig. 7**) suggests that it resides in the vicinity of proteins in the thylakoid membranes that are associated with PSI or PSII. A number of the proteins in its locality are members of the light harvesting family, both PSI and PSII types. Therefore, while enriched around the pyrenoid, RMT2 would still be present in the stroma where it could interact with proteins in both the stroma lamellae and grana stacks. These results raise the possibility that the function of RMT2 is associated with the proper assembly and interactions of light harvesting components with reaction centers and that aberrations in this process causes less efficient PSI biogenesis, but this is speculative and requires further verifications.

Together, our high-throughput O_2_ sensitivity screen led to the discovery of the novel protein RMT2, which is required for normal accumulation of PSI. This function of RMT2 is supported by the findings that the levels of PSI are markedly reduced in the *rmt2* mutant, RMT2 potentially interacts with other PSI assembly factors (e.g. PSBP4) and also with PsaB, and *rmt2* mutants displayed elevated levels of Ycf3 and Ycf4, two proteins previously shown to be critical for PSI assembly (11). Based on homology, RMT2 may be involved in protein methylation, even though it doesn’t have all features of known methyltransferases. The work also raises the possibility of a potential role of pyrenoids for compartmentation of O_2_-sensitive processes such as PSI assembly, although this function may be modulated by chloroplast oxidative conditions and only become prominent as the probability of oxidation of assembly intermediates increases. To understand RMT2 function, it is critical to determine the precise role of RMT2 in PSI assembly and whether it has an activity associated with protein methylation. It is also critical to obtain further biochemical results that define the protein interaction network of RMT2, if that network is enriched in the pyrenoid, and how it might integrate into assembly processes associated with photosynthetic complexes, especially during oxidative stress.

## Materials and Methods

### Strains and culture conditions

CMJ030 (CC-5325) was used as the WT strain for all experiments. Mutants in this study include *rmt2-1* (LMJ.RY0402.237082) and *rmt2-2* (LMJ.RY0402.255338), strains present in the CLiP mutant library (https://www.chlamylibrary.org/). Cells were grown in Tris Acetate Phosphate (TAP) solid and liquid medium. For photoautotrophic growth, Tris Phosphate (TP) medium (no acetate) was used. For all experiments, unless otherwise noted, strains were grown in constant LL (30 μmol photons m^−2^ s^−1^) and aerated by shaking. For growth under hypoxic conditions, cells were bubbled with a gas mixture (10% house air; 89.96% N_2_; 0.04% CO_2_) with constant stirring at 350 rpm at the indicated light intensities.

### Pooled mutant library screen

Chlamydomonas mutants from the CliP library were collected after growth on agar plates in LL and combined to generate a master pool of the entire library, as previously described (16, 17). Four aliquots of the master pool were inoculated into four cultures of 1 L of TAP medium to a final density of 20,000 cells mL^−1^. The four cultures were grown under four different conditions: low light hypoxic (LL-Hy), low light oxic (LL-Ox), high light hypoxic (HL-Hy), and high light oxic (HL-Ox). Growth was maintained for each of the cultures until the cell density reached approximately 2.5 million cells mL^−1^ (7 doublings on average). Following the various treatments, total genomic DNA was extracted and isolated from the cells and used to amplify the unique barcodes associated with individual mutants (16); the amplification products were sequenced to quantify the level of each mutant in the culture.

### Analysis of O_2_ sensitivity screen

Based on barcode sequencing results from 3 replicates for each of the experimental treatments, each barcode in the sample was assigned read counts. Several criteria were sequentially applied to filter the initial counts including: insertions in the 5’ UTR, CDS, or intron regions of the gene; an insertion confidence level, defined in the initial analysis of the CLiP library and recorded in the description of each mutant (https://www.chlamylibrary.org/allMutants), of 4 or better (from 53% to 95% confidence), the mean of the initial counts from three replicate experiments (using different pooled cultures) and from the final oxic HL and hypoxic HL conditions were each required to be greater than 10. Additionally, 2 or more mutant alleles per gene were required for the mutants to be further considered. The O_2_ tolerance for the mutants was calculated according to (16) with a threshold for defining a mutant as having an O_2_-sensitive phenotype set at 10 times higher counts+1 in hypoxic high light vs oxic high light conditions. For each gene, a p-value was generated using Fisher’s exact test comparing the numbers of alleles in that gene with and without a phenotype to the numbers of all insertions in the screen with and without the selected phenotype. A false discovery rate correction of the p-values was performed using the Benjamini-Hochberg (BH) procedure. For the purposes of displaying the fold-change in abundance in cells transferred from oxic to hypoxic conditions (Hy/Ox ratio), the read counts of each specific gene were averaged for hypoxic conditions and then divided by the average for oxic conditions. The Hy/Ox ratios were interpreted as a quantification of the average differential fitness of each mutant associated with a specific mutant allele, under hypoxic compared to oxic conditions. The p-value of the read counts was calculated using a parametric bootstrap and modeled as Poisson counts.

### Chlorophyll fluorescence analysis

Chl fluorescence to evaluate photosynthetic electron transport and ΦPSII was measured using a DUAL PAM-100 fluorometer. Cells were adapted in the dark for 20 min. Light curves were generated by illuminating cells with increasing light intensities (0, 10, 50, 100, 200, 400, 800, 1000, 1200 μmol photons m^−2^ s^−1^) with 1 min of illumination at each intensity. 1 mM NaHCO_3_ was added to the reaction mixture as an electron acceptor through CO_2_ fixation by the Calvin-Benson Cycle. Additional details are described in the legend of **Fig. 3**.

### P700 activity measurements

Absorbance changes associated with P700 oxidation and reduction were monitored at 705 nm and 740 nm using a JTS-100 spectrophotometer (SpectroLogiX, TN). Cells were pelleted by centrifugation and resuspended to a concentration of 30 μg Chl mL^−1^ in 20 mM HEPES-KOH, pH 7.2 and 10% ficoll. The suspension was made 10 μM DCMU to inhibit LEF prior to the measurement. Dark-adapted cells were exposed to 165 μmol photons m^−2^ s^−1^ during PSI oxidation followed by a saturating pulse as described in previous studies (9).

### ^15^N-labeling procedures for Chlamydomonas

Single colonies (of WT, *rmt2-1* or *rmt2-2* mutant cells) grown on TAP agar plates were inoculated in 5 mL of ^15^N-TAP medium, in which ^14^NH_4_Cl, the sole nitrogen source, was replaced with ^15^NH_4_Cl. After 5 d of growth in TAP, LL oxic conditions, the cells were diluted 100-fold in 50 mL of fresh ^15^N-TAP medium and then allowed to grow under appropriate conditions for the remainder of the experiment (2 additional days). For reciprocal labeling of mutant and WT cells, we performed the experiment identically, but the cells were labeled with ^14^N-TAP.

### Protein extraction and SDS-PAGE

Cultures containing 30 μg of Chl were collected by centrifugation at 3000 x *g* for 5 min, lysed and solubilized in 1X SDS-sample buffer (Cat. #1610737; Bio-Rad), heated at 95 °C for 2 min and then centrifuged for 5 min to pellet cell debris. 10 μL of the supernatant containing 3 μg of Chl of a ^14^N-labeled sample (e.g. mutant cells) was mixed with 10 μL of the supernatant containing the same amount of a ^15^N-labeled sample (e.g. WT cells). Thus, a 20 μL mixture of ^14^N- and ^15^N-labeled samples was loaded onto a Tris-glycine 4-20% polyacrylamide gel (Mini-PROTEAN TGX; Bio-Rad) and the proteins of the sample separated by electrophoresis at 60 V for 1 h. Reciprocal labelling of WT and mutant cells were also performed.

### In-gel digestion for LC/MS-MS

Proteins were resolved by SDS-PAGE and different regions of the gel (based on molecular mass) were excised, diced into small pieces (∼1 mm^2^), and placed into 0.65 mL tubes. The pieces were washed 3X by adding 200 µL of 25 mM ammonium bicarbonate/50% acetonitrile (ACN) followed by vortexing the suspension for 10 min, collecting the supernatant, and reducing and alkylating the cysteine residues of the proteins in the supernatant with 10 mM dithiothreitol (DTT) and 50 mM iodoacetamide (IAM), respectively; DTT was added to the sample, the sample incubated at 56 °C for 1 h and then made 50 mM IAM and incubated at room temperature for 45 min. The gel pieces were washed with 25% ammonium bicarbonate/50% ACN, dried completely in a Speed Vac and treated with trypsin (5 ng mL^−1^) at 37 °C overnight. Peptides were extracted three times from the gel pieces by adding 100 µL of 0.1% formic acid/50% acetonitrile. The extracted peptides were dried in a Speed Vac, resuspended in 0.1% formic acid and then cleaned using a ziptip (Milipore ZipTips C18).

### Complementation of *rmt2* mutants

In most cases **t**he pRam118 plasmid with VENUS (68) was used for complementation of the *rmt2* mutant strains (in some cases mNeonGreen was used; see below). For the generation of the plasmid pRam118-RMT2-VENUS, genomic DNA for *RMT2* with an extended 5’ UTR was assembled into the pRam118_VENUS plasmid using Gibson assembly (69). The mutant strain *rmt2-2* was transformed with the plasmid pRam118-RMT2-VENUS and transformants screened for VENUS fluorescence using a microplate reader (Infinite M1000; TECAN) as previously described (68). Excitation and emission settings were, VENUS excitation 515/12 nm and emission 550/12 nm.

### Subcellular Localization of RMT2

The pJM43-ble plasmid, which contains the gene encoding Bleomycin resistance, with a mNeonGreen tag from pYF015, was used for expression of *RMT2* (16). For the generation of the pJM43-ble-RMT2-mNeonGreen construct expressing the *RMT2* gene, genomic *RMT2* DNA with the endogenous promoter and complete 5’ UTR was assembled with the pJM43-ble-mNeonGreen plasmid using Gibson assembly (69). The *rmt2-2* mutant was transformed with plasmid pJM43-ble-RMT2-mNeonGreen, transformants were selected for bleomycin resistance and grown in moderate light and then screened for mNeonGreen fluorescence as described (68). Briefly, putative transgenic cells placed in individual wells of a microtiter plate were screened for mNeonGreen fluorescence using a microplate reader (Infinite M1000; TECAN). Excitation and emission settings were excitation 488/12 nm and emission 520/5 nm. Strains exhibiting mNeonGreen fluorescence were used to localize the RMT2-mNeonGreen fusion protein by TCS SP8 confocal laser-scanning microscopy (Leica); excitation/emission settings were 488 nm/515-530 nm using the HyD SMD hybrid detector for mNeonGreen, and 488 nm/650-700 using the nm-HyD SMD hybrid detector for Chl autofluorescence.

### Electron microscopy

Cells were cultured in TAP medium in 20 µmol photons m^−2^ s^−1^ or in HS medium in 50 µmol photons m^−2^ s^−1^ and grown to a concentration of ∼5 µg Chl mL^−1^. The cells were harvested by filtration, fixed with 3 % glutaraldehyde and 4 % paraformaldehyde in 0.1 M sodium cacodylate buffer, pH 6.8, first for 1 h at room temperature, then at 4 °C overnight. The fixed cells were washed three times with the same buffer and then embedded in 2 % low-melting-temperature agarose (SeaPlaque, Lonza). The embedded samples were post-fixed with 4 % potassium permanganate at 4 °C for 2 h, washed three times with H_2_O, incubated in 2 % uranyl acetate at room temperature for 1 h, washed twice with H_2_O, and dehydrated through a graded series of increasing ethanol concentrations. Sections of ∼70 nm were cut using a Leica Ultracut S microtome, collected on formvar-coated 100-mesh copper grids (Electron Microscopy Sciences), and post-stained for 30 sec in 1:1 3 % uranyl acetate and 50% acetone, followed by 0.2 % lead citrate for 3 min. Sections were then imaged at 120 kV using a JEM-1400 transmission electron microscope (JEOL) equipped with a Gatan Orius 4k X 4k digital camera.

### PSI Turnover

CMJ030, *rmt2-2* and *rmt2-2* (RMT2-mNG) cells were grown in TAP medium to mid-log phase. The cultures were then centrifuged at 3000 x *g* for 5 min and cell pellets resuspended to 5 µg Chl mL^−1^. 100 mL of cell suspension was transferred to two 250 mL Erlenmeyer flasks and one flask of each strain was treated simultaneously with chloramphenicol (250 µg mL^−1^) and cycloheximide (10 µg mL^−1^), which inhibit translation on 70S and 80S ribosomes, respectively. Cells of control and treated samples were harvested after 0, 6 and 24 h and proteins were extracted as described above.

### Proximity labeling of RMT2

A DNA construct containing the native RMT2 promoter and the PSAD terminator controlling the gene encoding RMT2-BioID fusion protein linked to mNeonGreen (RMT2-promoter::RMT2-BioID-G3-mNeonGreen::PSAD-terminator) was transformed into the *rmt2*-*1* and *rmt2-2* mutants. Controls include RMT2-VENUS in the *rmt2-1* and *rmt2-2* mutant background. The strains were labeled during growth in LL (30 μmol photons m^−2^ s^−1^) in TAP-^14^N or TAP-^15^N liquid medium, as described above, and treated with 400 μM biotin for 4 h before being harvested. Proteins were extracted, digested to peptides, and mixed. Biotinylated peptides were enriched as described (35) with 5% ACN used throughout the enrichment process.

### Mass Spectrometry and data analysis

LC-MS/MS was carried out on a Q-Exactive HF hybrid quadrupole-Orbitrap mass spectrometer (Thermo Fisher), equipped with an Easy LC 1200 UPLC liquid chromatography system (Thermo Fisher). Peptides were separated on a 50 cm analytical column ES803 (ThermoFisher). The flow rate was 300 nL min^−1^, and a 120-min gradient was used. Peptides were eluted from the column using the gradients; from 3-28% solvent B (80% acetonitrile, 0.1 % formic acid) over 100 min and then from 28-44% solvent B over 20 min, followed by a short wash with 90% solvent B. Precursor scan was from mass-to-charge ratio (m/z) 375 to 1,600 and the top 20 most intense multiply charged precursors were selected for fragmentation. Peptides were fragmented with higher-energy collision dissociation (HCD) with normalized collision energy (NCE) 27.

MS/MS spectra were searched by Protein Prospector as described in (70), against a database constructed from Chlamydomonas proteins (Phytozome v5.6 and 6.1) with a decoy database of reversed peptides. Precursor and fragment tolerances were set to 10 ppm and 20 ppm, respectively. Biotinylated peptides were searched using exogenous biotin (ExBio) on lysine as a variable modification. Total protein quantification was performed as described in (70). For quantification of biotinylated peptides, unmodified peptides were first filtered out (quantification shows 1:1 ratio, showing equal mixture) and the median number of the ratio of biotinylated peptides (RMT2-ID/control) for each protein was calculated from each experiment. The fold of 100 was used for all ratios that are greater than 100-fold difference, converted to log2 numbers, and then the median was calculated. For a few biotinylated peptides with missing identifications/quantifications in replicates, MS1 peaks were manually examined using Skyline (71) at the specified retention time for ratio calculation based on similarly detected peptides. The diagnostic ion peak of 310.16 in ^14^N-labeled samples and 311.16 in ^15^N-labeled biotinylated peptides, corresponding to the derivative ion due to ammonia loss from the immonium ion of biotinylated lysine (ImKBio), was manually examined for quality control.

## Acknowledgments

RGK and JF were supported by the Department of Plant Biology of the Carnegie Institution for Science. WH was supported by the Department of Energy (DOE) Grant DE-SC0019417 (to ARG). PR was supported by the Human Frontier of Science Program (HFSP) RGP0046/2018 (to ARG). FB was supported by the National Science Foundation (NSF) grant 1921429 (to ARG and Devaki Bhaya). MO was supported by NSF Grant MCB 1818383. RS was supported by the National Institutes of Health grant R01GM135706 (to S-LX). Andrey Malkovskiy provided help with the confocal microscopy while RS and TG were major contributors to the mass-spec analysis, which was performed at the Carnegie MS facility.

