## Supplemental figures and tables for "Chloroplast Methyltransferase Homolog RMT2 is Involved in Photosystem I Biogenesis"

### Slide 1
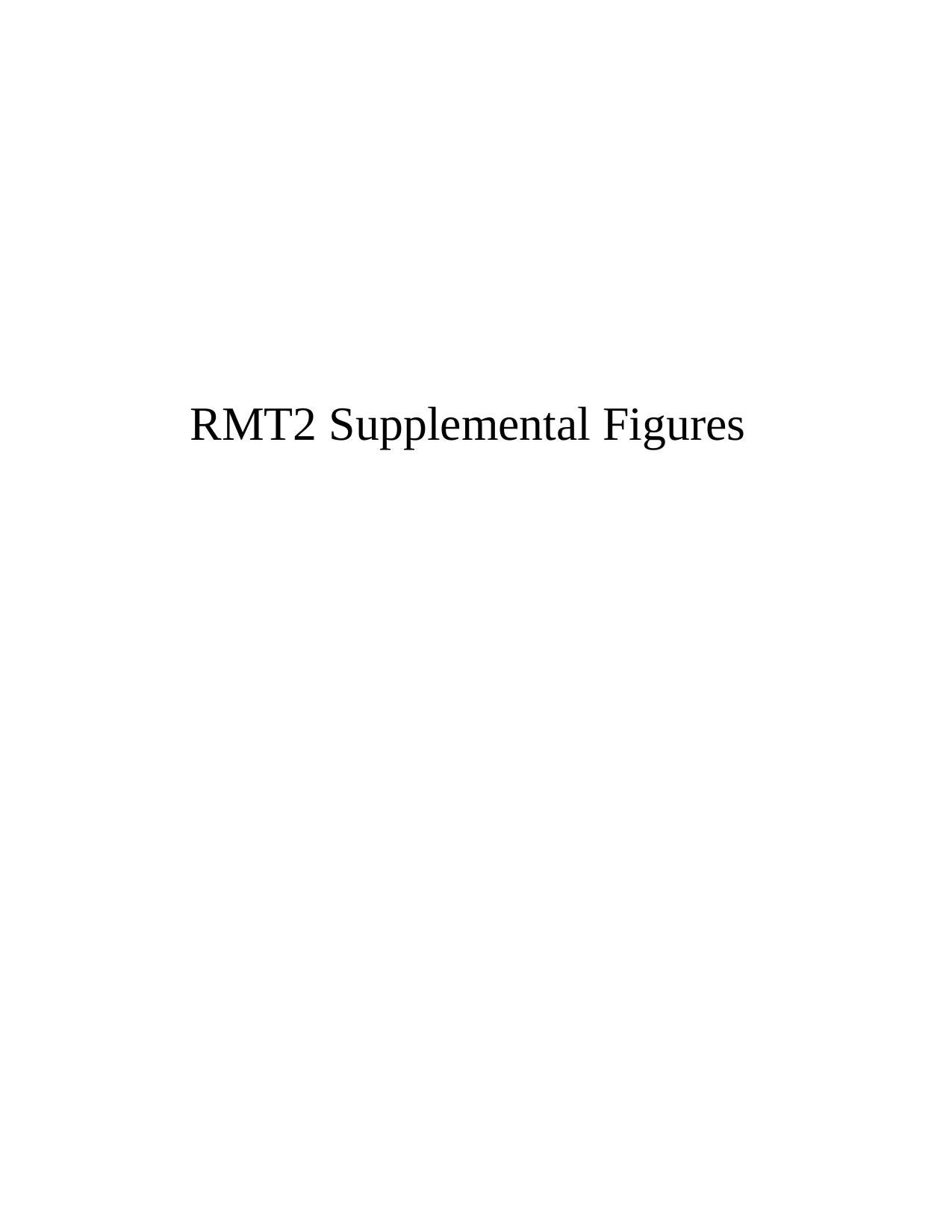

#### RMT2 Supplemental Figures

### Slide 2
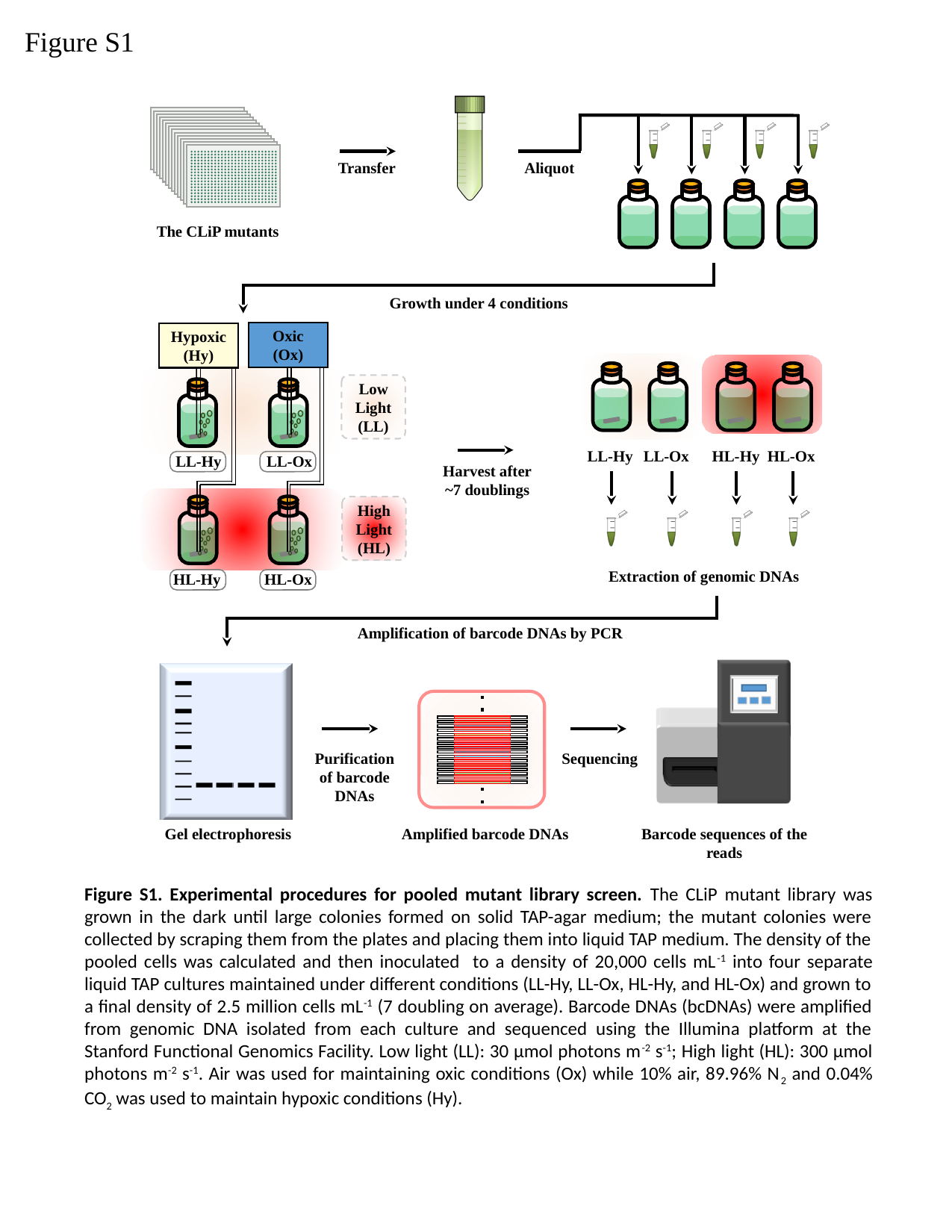

Figure S1
Transfer
Aliquot
The CLiP mutants
Growth under 4 conditions
Oxic
(Ox)
Hypoxic
(Hy)
Low Light
(LL)
LL-Hy
LL-Ox
HL-Hy
HL-Ox
LL-Hy
LL-Ox
Harvest after ~7 doublings
High Light
(HL)
Extraction of genomic DNAs
HL-Hy
HL-Ox
Amplification of barcode DNAs by PCR
Purification of barcode DNAs
Sequencing
Barcode sequences of the reads
Gel electrophoresis
Amplified barcode DNAs
Figure S1. Experimental procedures for pooled mutant library screen. The CLiP mutant library was grown in the dark until large colonies formed on solid TAP-agar medium; the mutant colonies were collected by scraping them from the plates and placing them into liquid TAP medium. The density of the pooled cells was calculated and then inoculated to a density of 20,000 cells mL^-1^ into four separate liquid TAP cultures maintained under different conditions (LL-Hy, LL-Ox, HL-Hy, and HL-Ox) and grown to a final density of 2.5 million cells mL^-1^ (7 doubling on average). Barcode DNAs (bcDNAs) were amplified from genomic DNA isolated from each culture and sequenced using the Illumina platform at the Stanford Functional Genomics Facility. Low light (LL): 30 µmol photons m^-2^ s^-1^; High light (HL): 300 µmol photons m^-2^ s^-1^. Air was used for maintaining oxic conditions (Ox) while 10% air, 89.96% N_2_ and 0.04% CO_2_ was used to maintain hypoxic conditions (Hy).

### Slide 3
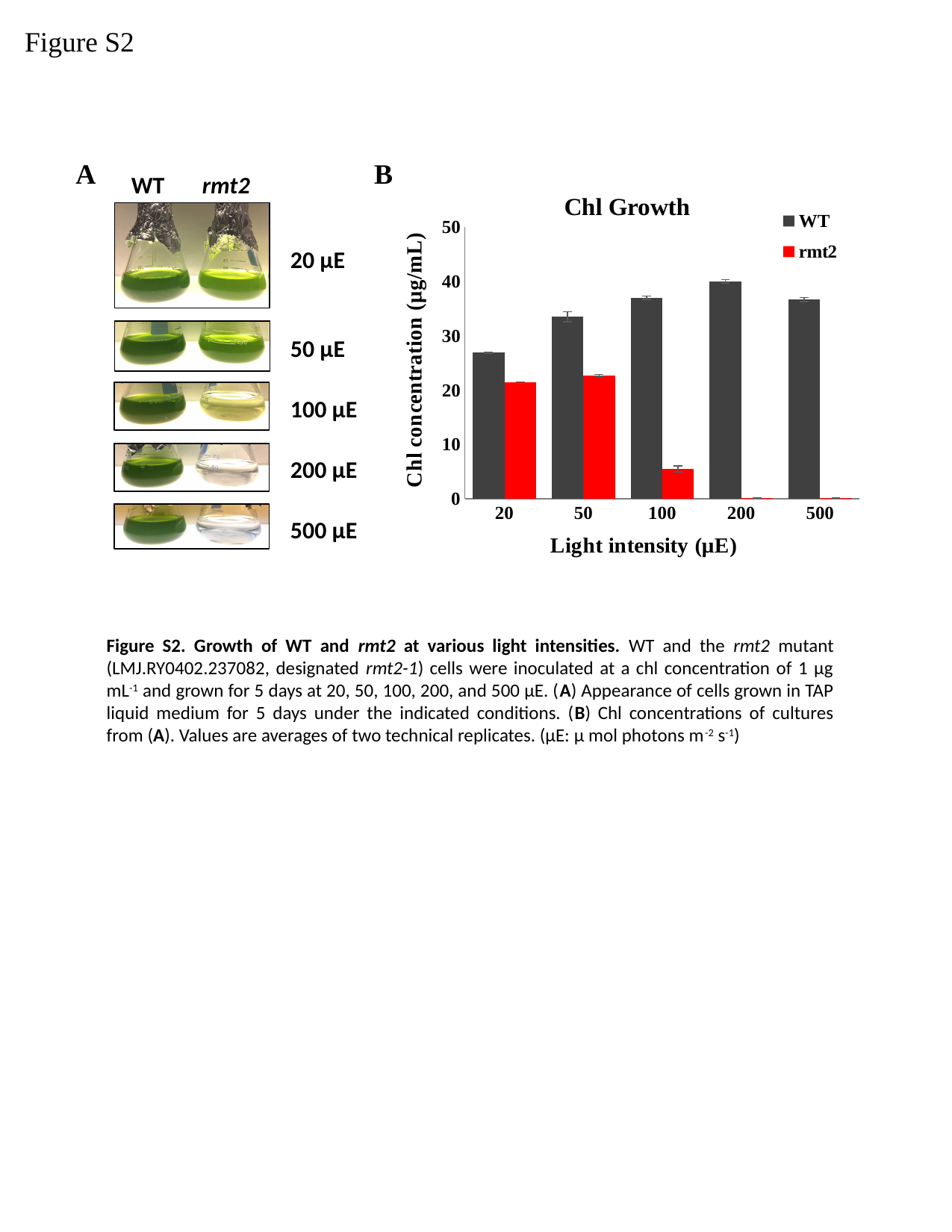

Figure S2
A
B
rmt2
WT
20 μE
50 μE
100 μE
200 μE
500 μE
###### Chart: Chl Growth
| Category | WT | rmt2 |
|---|---|---|
| 20 | 26.94 | 21.405 |
| 50 | 33.510000000000005 | 22.740000000000002 |
| 100 | 37.06 | 5.505 |
| 200 | 40.06 | 0.175 |
| 500 | 36.7 | 0.14500000000000002 |Figure S2. Growth of WT and rmt2 at various light intensities. WT and the rmt2 mutant (LMJ.RY0402.237082, designated rmt2-1) cells were inoculated at a chl concentration of 1 μg mL^-1^ and grown for 5 days at 20, 50, 100, 200, and 500 μE. (A) Appearance of cells grown in TAP liquid medium for 5 days under the indicated conditions. (B) Chl concentrations of cultures from (A). Values are averages of two technical replicates. (µE: µ mol photons m^-2^ s^-1^)

### Slide 4
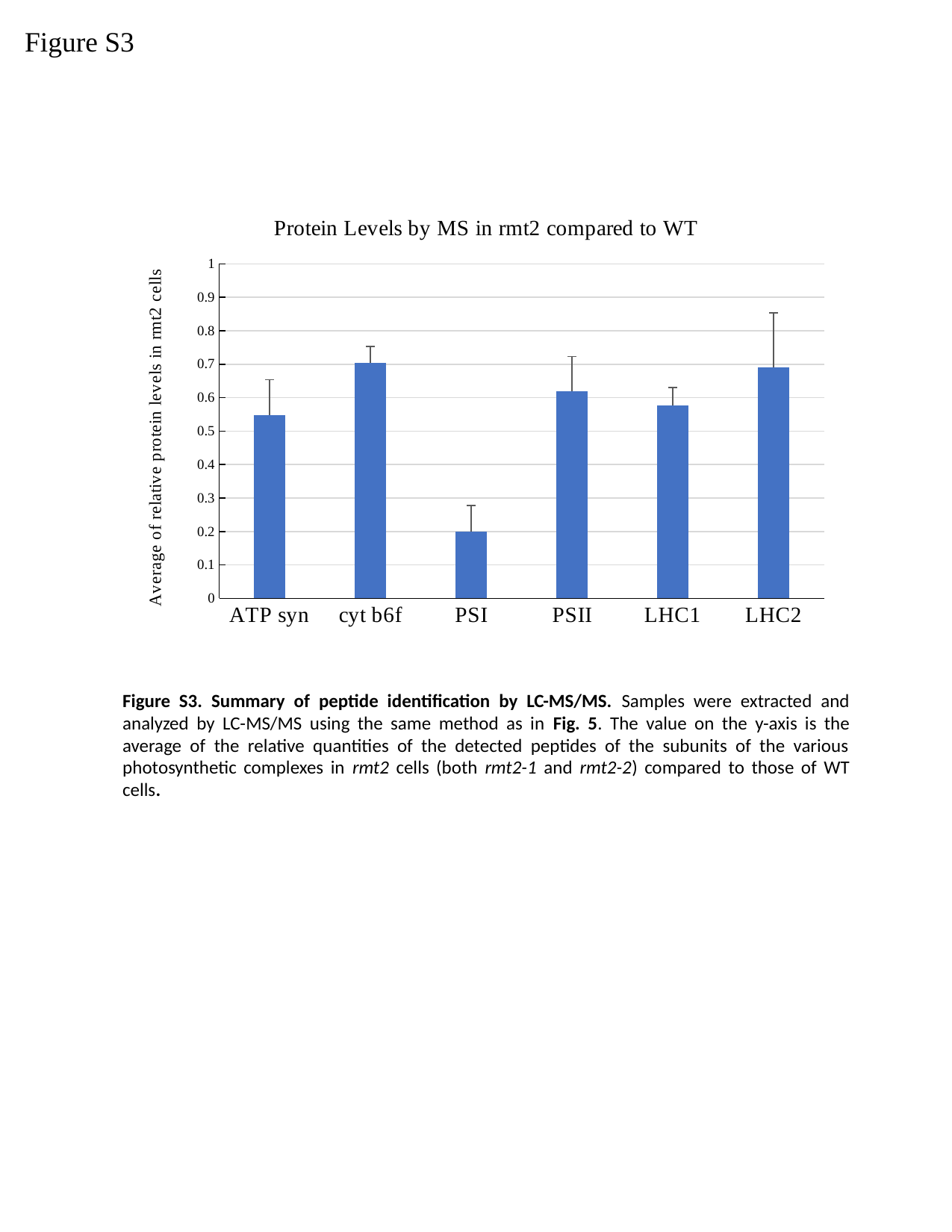

Figure S3
###### Chart: Protein Levels by MS in rmt2 compared to WT
| Category | |
|---|---|
| ATP syn | 0.548448179321665 |
| cyt b6f | 0.7048375007929681 |
| PSI | 0.20014806533320725 |
| PSII | 0.6196727408501059 |
| LHC1 | 0.57667598875759 |
| LHC2 | 0.6895622003530677 |Figure S3. Summary of peptide identification by LC-MS/MS. Samples were extracted and analyzed by LC-MS/MS using the same method as in Fig. 5. The value on the y-axis is the average of the relative quantities of the detected peptides of the subunits of the various photosynthetic complexes in rmt2 cells (both rmt2-1 and rmt2-2) compared to those of WT cells.

### Slide 5
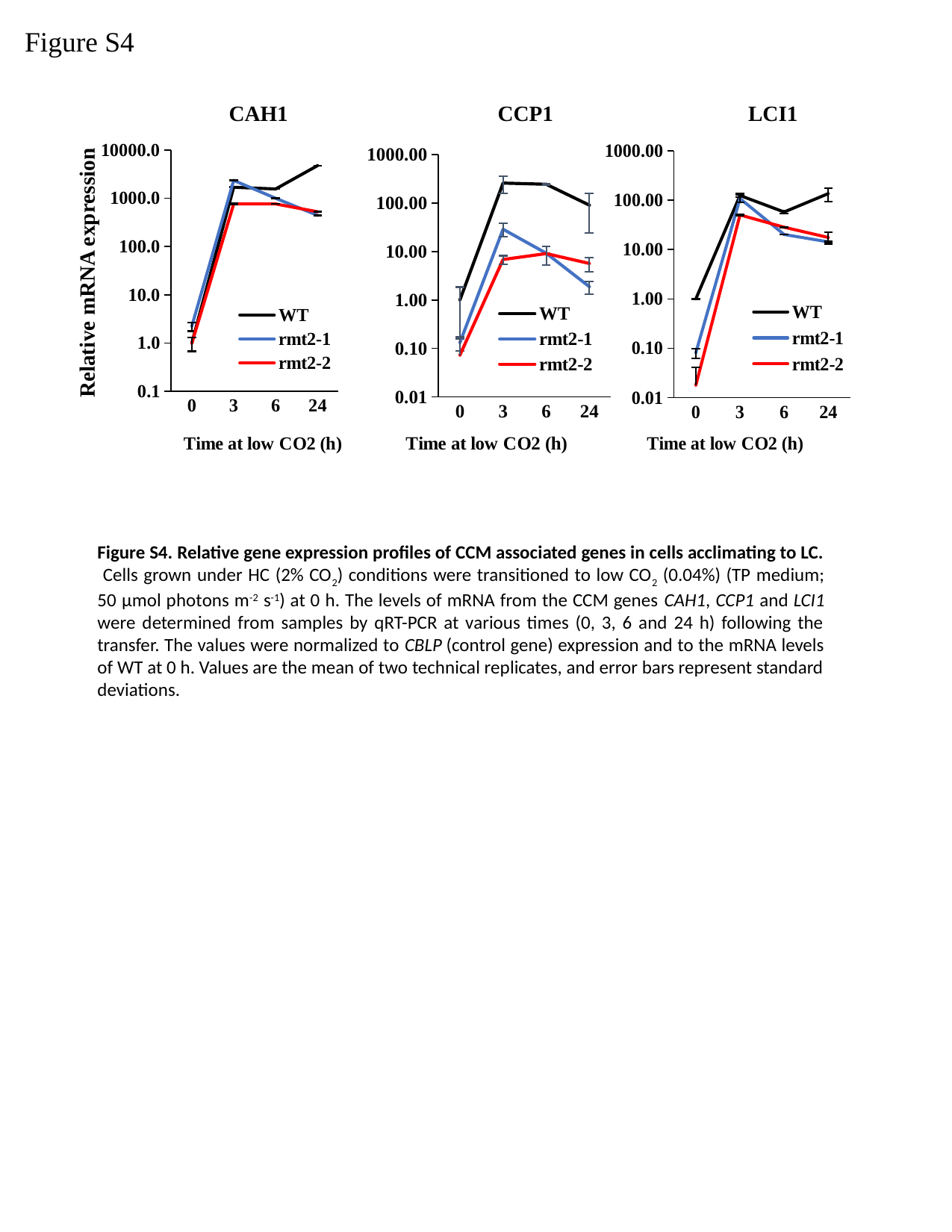

Figure S4
CAH1
CCP1
LCI1
###### Chart
| Category | | | |
|---|---|---|---|
| 0 | 1.0 | 0.08122783366449192 | 0.017878915861216385 |
| 3 | 124.29864870180994 | 107.6682645205998 | 50.23298398635828 |
| 6 | 57.524661360777856 | 20.11028155232479 | 28.245308493202653 |
| 24 | 135.57185689735863 | 14.276427202996231 | 17.470680018800863 |
###### Chart
| Category | | | |
|---|---|---|---|
| 0 | 1.0 | 2.233796304883136 | 0.9876596427952106 |
| 3 | 1685.2569030069808 | 2342.369274604469 | 762.3990346515095 |
| 6 | 1555.9121459777534 | 995.084235235862 | 761.9596964739696 |
| 24 | 4733.103906538219 | 442.20480114998327 | 524.1544280058185 |
###### Chart
| Category | | | |
|---|---|---|---|
| 0 | 1.0 | 0.13209209129036942 | 0.07262668111782677 |
| 3 | 257.52303099744364 | 28.765512083706273 | 6.844215181835434 |
| 6 | 243.3296053329583 | 9.196878811148313 | 9.045171888390803 |
| 24 | 90.2038605408483 | 1.87518864942455 | 5.670530137862575 |Relative mRNA expression
Figure S4. Relative gene expression profiles of CCM associated genes in cells acclimating to LC. Cells grown under HC (2% CO_2_) conditions were transitioned to low CO_2_ (0.04%) (TP medium; 50 µmol photons m^-2^ s^-1^) at 0 h. The levels of mRNA from the CCM genes CAH1, CCP1 and LCI1 were determined from samples by qRT-PCR at various times (0, 3, 6 and 24 h) following the transfer. The values were normalized to CBLP (control gene) expression and to the mRNA levels of WT at 0 h. Values are the mean of two technical replicates, and error bars represent standard deviations.

### Slide 6
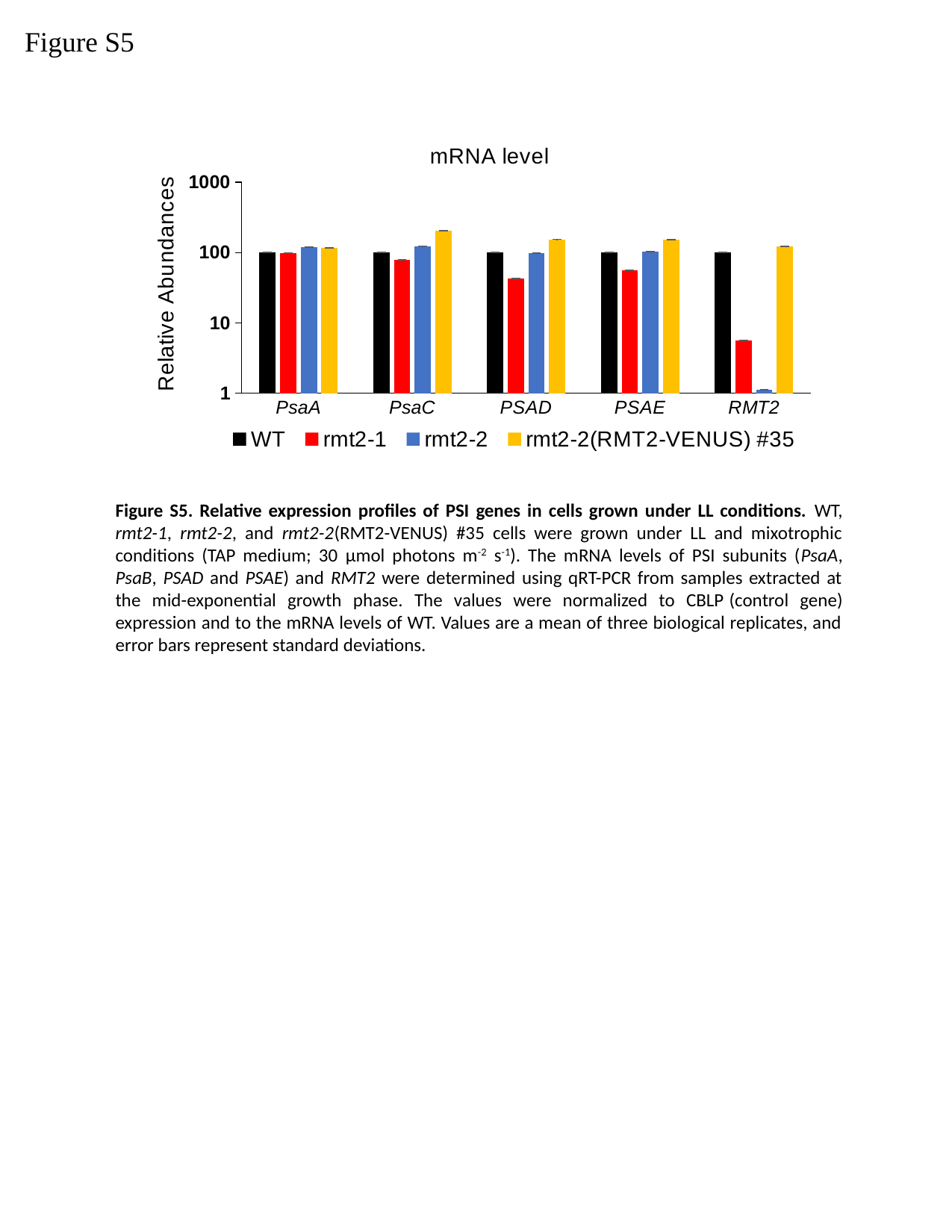

Figure S5
###### Chart: mRNA level
| Category | WT | rmt2-1 | rmt2-2 | rmt2-2(RMT2-VENUS) #35 |
|---|---|---|---|---|
| PsaA | 100.0 | 98.76604552365636 | 120.63783800587679 | 116.66676549466548 |
| PsaC | 100.0 | 79.02551659408952 | 123.69525948690475 | 203.02892447992454 |
| PSAD | 100.0 | 42.83329299717409 | 98.41482182168407 | 154.21464100548923 |
| PSAE | 100.0 | 56.051924921717465 | 103.91657072564131 | 152.51147193060368 |
| RMT2 | 100.0 | 5.66783236036367 | 1.12906584714103 | 121.97664374416 |Figure S5. Relative expression profiles of PSI genes in cells grown under LL conditions. WT, rmt2-1, rmt2-2, and rmt2-2(RMT2-VENUS) #35 cells were grown under LL and mixotrophic conditions (TAP medium; 30 µmol photons m^-2^ s^-1^). The mRNA levels of PSI subunits (PsaA, PsaB, PSAD and PSAE) and RMT2 were determined using qRT-PCR from samples extracted at the mid-exponential growth phase. The values were normalized to CBLP (control gene) expression and to the mRNA levels of WT. Values are a mean of three biological replicates, and error bars represent standard deviations.

### Slide 7
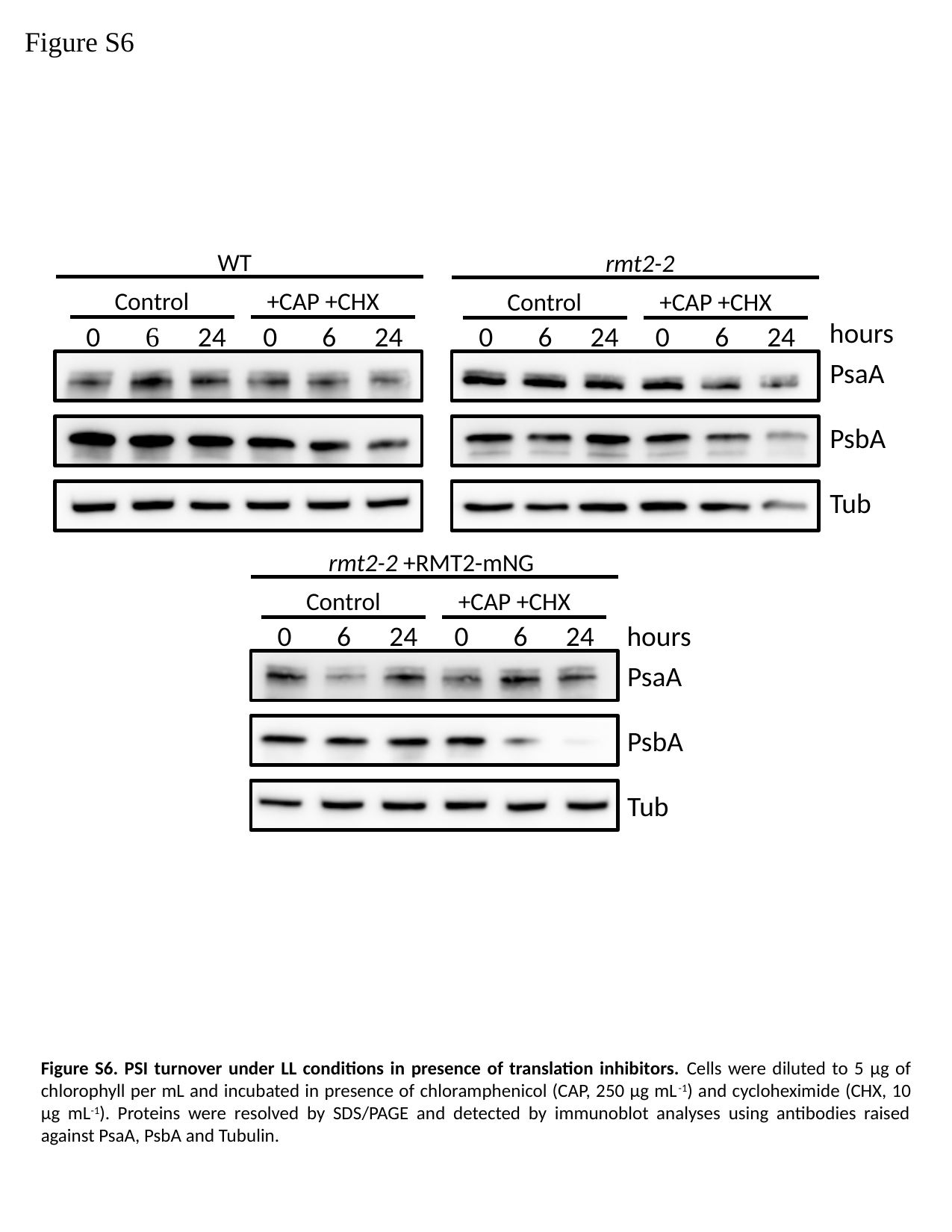

Figure S6
WT
rmt2-2
Control
+CAP +CHX
Control
+CAP +CHX
hours
24
24
0
6
0
6
24
24
0
6
0
6
PsaA
PsbA
Tub
rmt2-2 +RMT2-mNG
Control
+CAP +CHX
hours
0
6
0
6
24
24
PsaA
PsbA
Tub
Figure S6. PSI turnover under LL conditions in presence of translation inhibitors. Cells were diluted to 5 µg of chlorophyll per mL and incubated in presence of chloramphenicol (CAP, 250 µg mL^-1^) and cycloheximide (CHX, 10 µg mL^-1^). Proteins were resolved by SDS/PAGE and detected by immunoblot analyses using antibodies raised against PsaA, PsbA and Tubulin.

### Slide 8
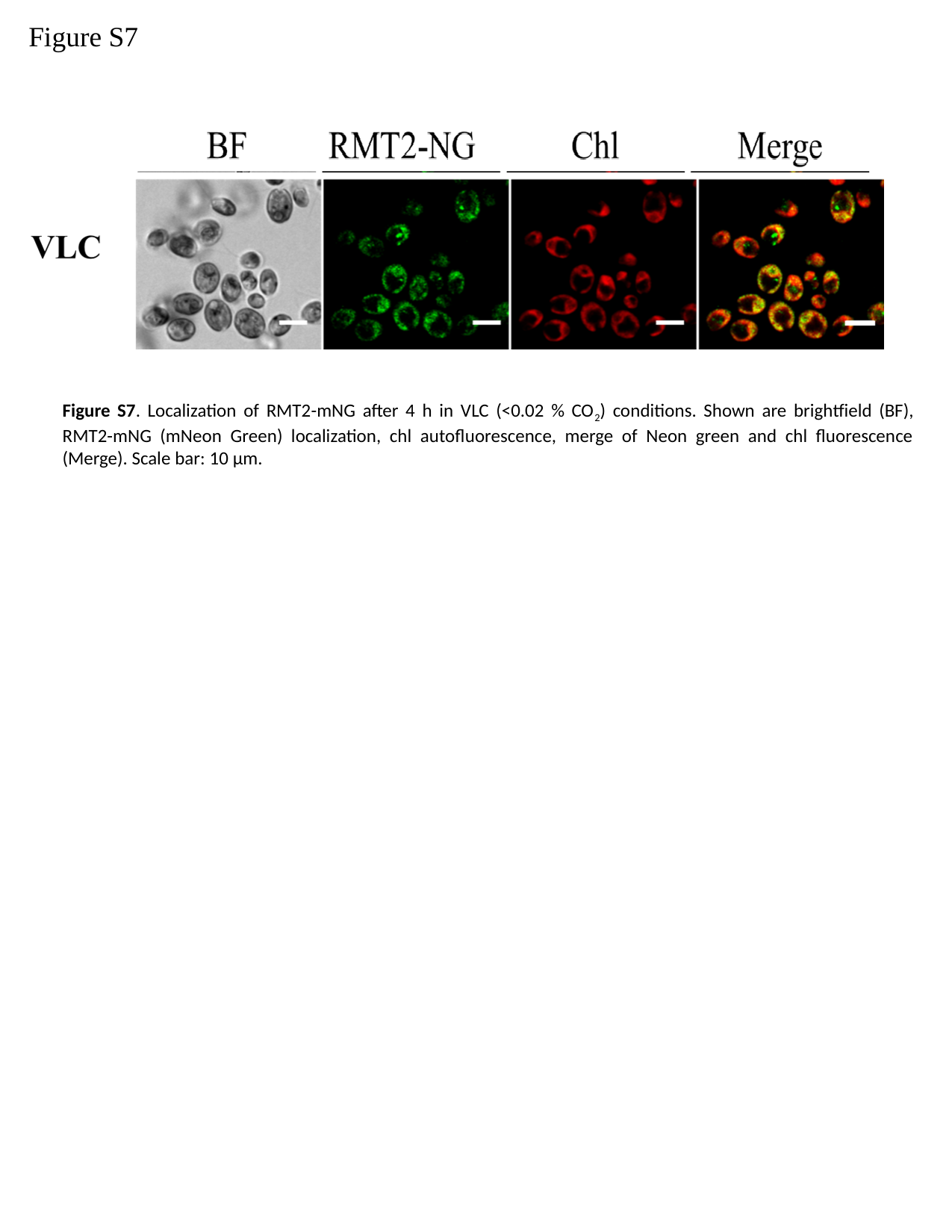

Figure S7
Figure S7. Localization of RMT2-mNG after 4 h in VLC (<0.02 % CO_2_) conditions. Shown are brightfield (BF), RMT2-mNG (mNeon Green) localization, chl autofluorescence, merge of Neon green and chl fluorescence (Merge). Scale bar: 10 µm.

### Slide 9
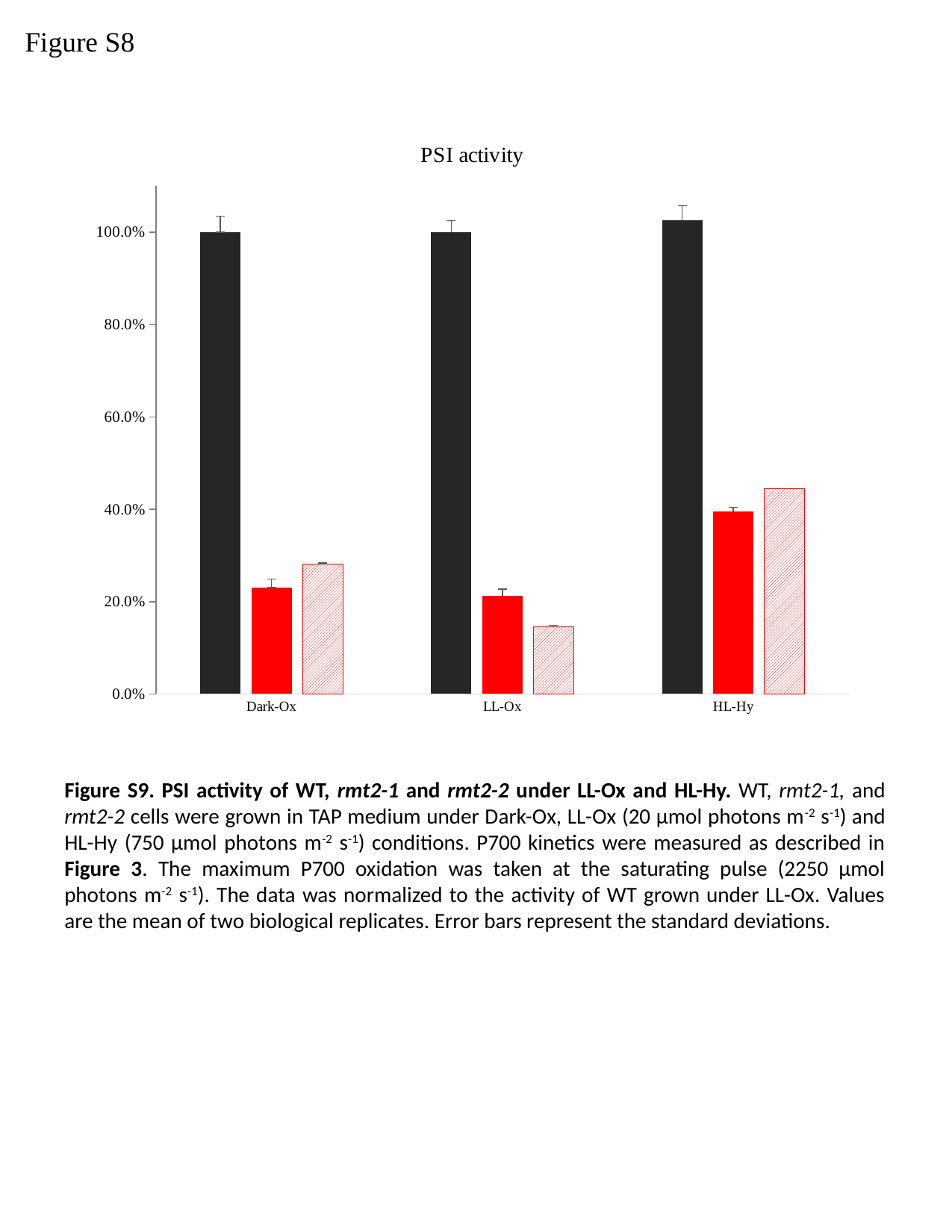

Figure S8
###### Chart: PSI activity
| Category | WT | rmt2-1 | rmt2-2 |
|---|---|---|---|
| Dark-Ox | 1.0 | 0.23011403254211998 | 0.28195768180370917 |
| LL-Ox | 1.0 | 0.21253564885108817 | 0.14565178837861742 |
| HL-Hy | 1.0257903103708 | 0.3958386769446429 | 0.44509882094461245 |Figure S9. PSI activity of WT, rmt2-1 and rmt2-2 under LL-Ox and HL-Hy. WT, rmt2-1, and rmt2-2 cells were grown in TAP medium under Dark-Ox, LL-Ox (20 µmol photons m^-2^ s^-1^) and HL-Hy (750 µmol photons m^-2^ s^-1^) conditions. P700 kinetics were measured as described in Figure 3. The maximum P700 oxidation was taken at the saturating pulse (2250 µmol photons m^-2^ s^-1^). The data was normalized to the activity of WT grown under LL-Ox. Values are the mean of two biological replicates. Error bars represent the standard deviations.

### Slide 10
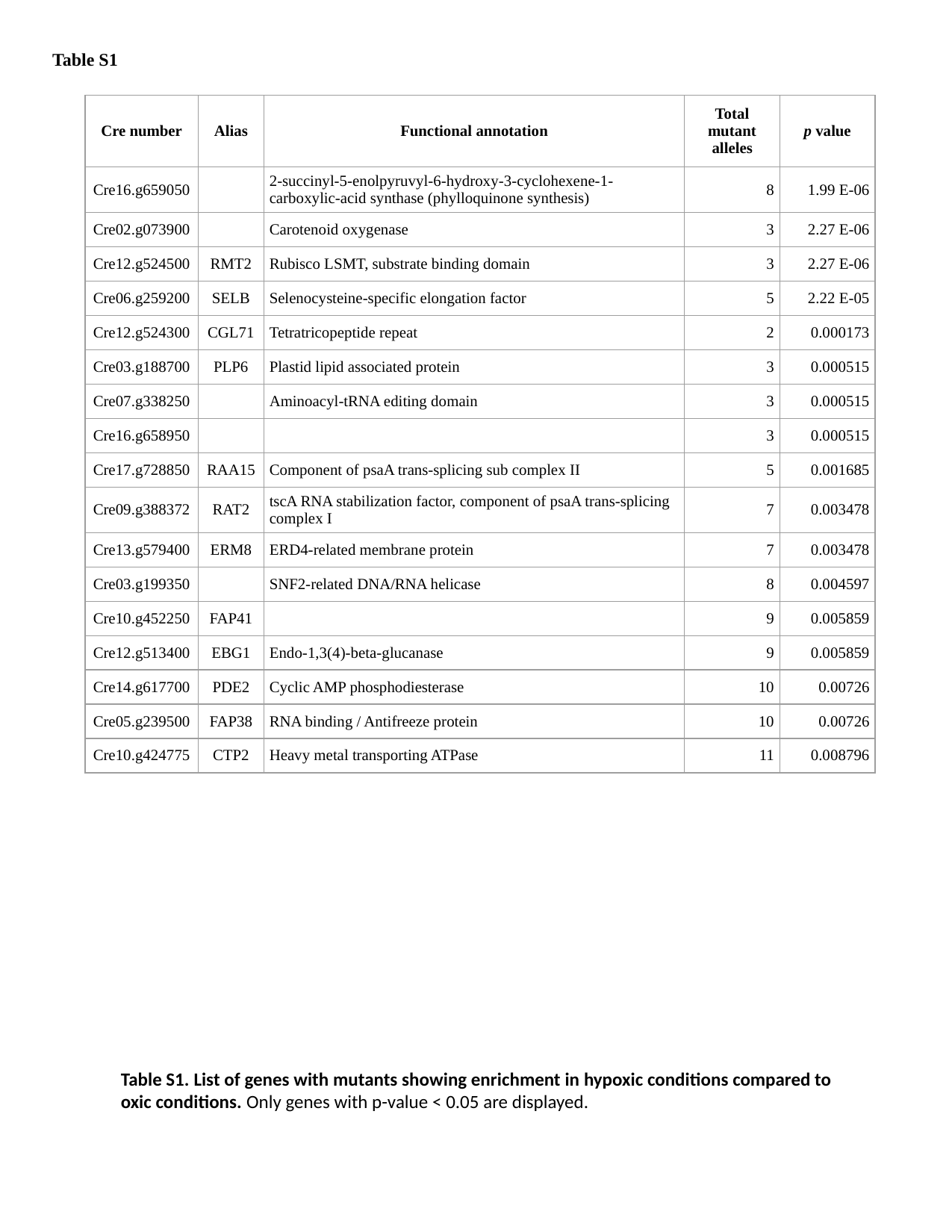

Table S1
| Cre number | Alias | Functional annotation | Total mutant alleles | p value |
| --- | --- | --- | --- | --- |
| Cre16.g659050 | | 2-succinyl-5-enolpyruvyl-6-hydroxy-3-cyclohexene-1-carboxylic-acid synthase (phylloquinone synthesis) | 8 | 1.99 E-06 |
| Cre02.g073900 | | Carotenoid oxygenase | 3 | 2.27 E-06 |
| Cre12.g524500 | RMT2 | Rubisco LSMT, substrate binding domain | 3 | 2.27 E-06 |
| Cre06.g259200 | SELB | Selenocysteine-specific elongation factor | 5 | 2.22 E-05 |
| Cre12.g524300 | CGL71 | Tetratricopeptide repeat | 2 | 0.000173 |
| Cre03.g188700 | PLP6 | Plastid lipid associated protein | 3 | 0.000515 |
| Cre07.g338250 | | Aminoacyl-tRNA editing domain | 3 | 0.000515 |
| Cre16.g658950 | | | 3 | 0.000515 |
| Cre17.g728850 | RAA15 | Component of psaA trans-splicing sub complex II | 5 | 0.001685 |
| Cre09.g388372 | RAT2 | tscA RNA stabilization factor, component of psaA trans-splicing complex I | 7 | 0.003478 |
| Cre13.g579400 | ERM8 | ERD4-related membrane protein | 7 | 0.003478 |
| Cre03.g199350 | | SNF2-related DNA/RNA helicase | 8 | 0.004597 |
| Cre10.g452250 | FAP41 | | 9 | 0.005859 |
| Cre12.g513400 | EBG1 | Endo-1,3(4)-beta-glucanase | 9 | 0.005859 |
| Cre14.g617700 | PDE2 | Cyclic AMP phosphodiesterase | 10 | 0.00726 |
| Cre05.g239500 | FAP38 | RNA binding / Antifreeze protein | 10 | 0.00726 |
| Cre10.g424775 | CTP2 | Heavy metal transporting ATPase | 11 | 0.008796 |
Table S1. List of genes with mutants showing enrichment in hypoxic conditions compared to oxic conditions. Only genes with p-value < 0.05 are displayed.

### Slide 11
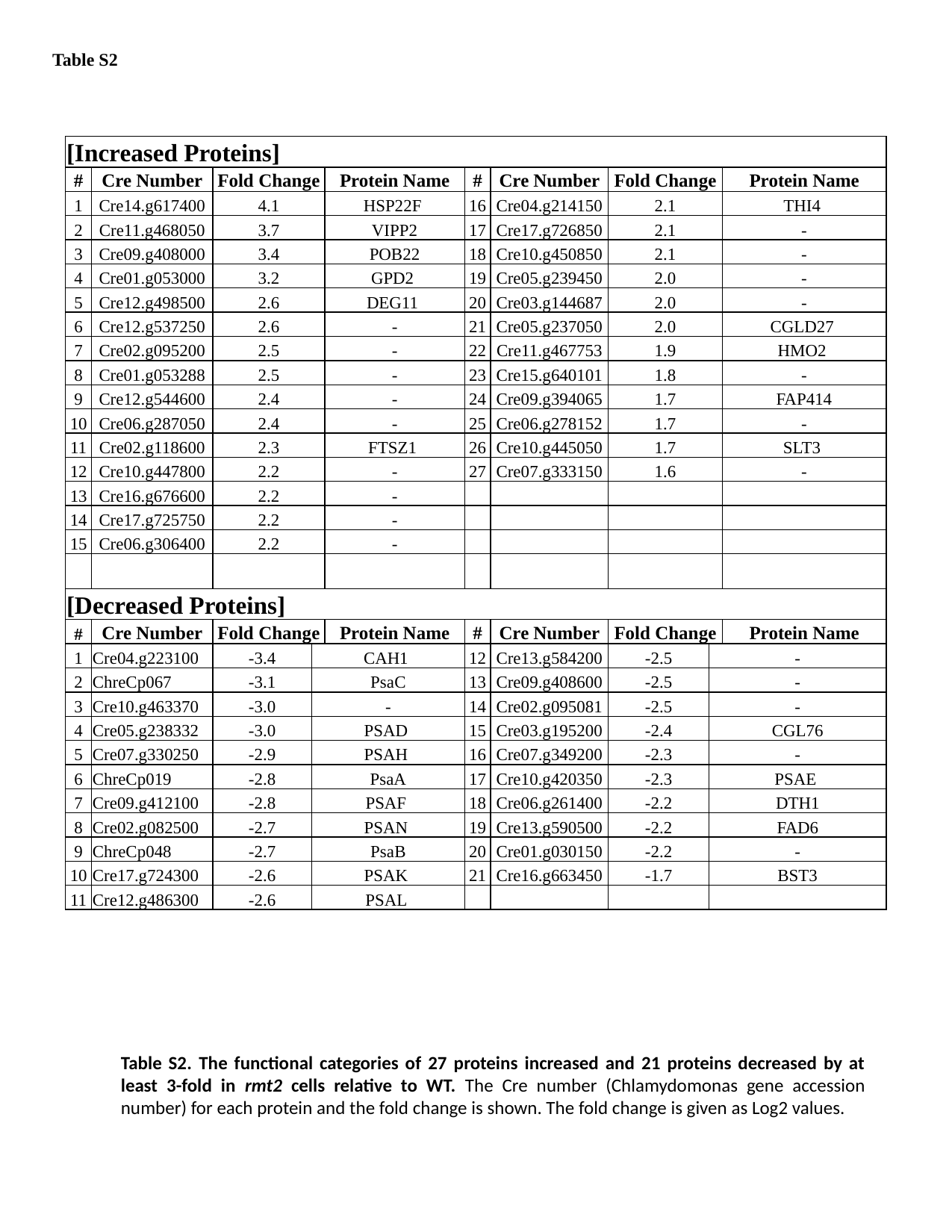

Table S2
| [Increased Proteins] | | | | | | | | | |
| --- | --- | --- | --- | --- | --- | --- | --- | --- | --- |
| # | Cre Number | Fold Change | | Protein Name | # | Cre Number | Fold Change | | Protein Name |
| 1 | Cre14.g617400 | 4.1 | | HSP22F | 16 | Cre04.g214150 | 2.1 | | THI4 |
| 2 | Cre11.g468050 | 3.7 | | VIPP2 | 17 | Cre17.g726850 | 2.1 | | - |
| 3 | Cre09.g408000 | 3.4 | | POB22 | 18 | Cre10.g450850 | 2.1 | | - |
| 4 | Cre01.g053000 | 3.2 | | GPD2 | 19 | Cre05.g239450 | 2.0 | | - |
| 5 | Cre12.g498500 | 2.6 | | DEG11 | 20 | Cre03.g144687 | 2.0 | | - |
| 6 | Cre12.g537250 | 2.6 | | - | 21 | Cre05.g237050 | 2.0 | | CGLD27 |
| 7 | Cre02.g095200 | 2.5 | | - | 22 | Cre11.g467753 | 1.9 | | HMO2 |
| 8 | Cre01.g053288 | 2.5 | | - | 23 | Cre15.g640101 | 1.8 | | - |
| 9 | Cre12.g544600 | 2.4 | | - | 24 | Cre09.g394065 | 1.7 | | FAP414 |
| 10 | Cre06.g287050 | 2.4 | | - | 25 | Cre06.g278152 | 1.7 | | - |
| 11 | Cre02.g118600 | 2.3 | | FTSZ1 | 26 | Cre10.g445050 | 1.7 | | SLT3 |
| 12 | Cre10.g447800 | 2.2 | | - | 27 | Cre07.g333150 | 1.6 | | - |
| 13 | Cre16.g676600 | 2.2 | | - | | | | | |
| 14 | Cre17.g725750 | 2.2 | | - | | | | | |
| 15 | Cre06.g306400 | 2.2 | | - | | | | | |
| [Decreased Proteins] | | | | | | | | | |
| # | Cre Number | Fold Change | | Protein Name | # | Cre Number | Fold Change | | Protein Name |
| 1 | Cre04.g223100 | -3.4 | CAH1 | | 12 | Cre13.g584200 | -2.5 | - | |
| 2 | ChreCp067 | -3.1 | PsaC | | 13 | Cre09.g408600 | -2.5 | - | |
| 3 | Cre10.g463370 | -3.0 | - | | 14 | Cre02.g095081 | -2.5 | - | |
| 4 | Cre05.g238332 | -3.0 | PSAD | | 15 | Cre03.g195200 | -2.4 | CGL76 | |
| 5 | Cre07.g330250 | -2.9 | PSAH | | 16 | Cre07.g349200 | -2.3 | - | |
| 6 | ChreCp019 | -2.8 | PsaA | | 17 | Cre10.g420350 | -2.3 | PSAE | |
| 7 | Cre09.g412100 | -2.8 | PSAF | | 18 | Cre06.g261400 | -2.2 | DTH1 | |
| 8 | Cre02.g082500 | -2.7 | PSAN | | 19 | Cre13.g590500 | -2.2 | FAD6 | |
| 9 | ChreCp048 | -2.7 | PsaB | | 20 | Cre01.g030150 | -2.2 | - | |
| 10 | Cre17.g724300 | -2.6 | PSAK | | 21 | Cre16.g663450 | -1.7 | BST3 | |
| 11 | Cre12.g486300 | -2.6 | PSAL | | | | | | |
Table S2. The functional categories of 27 proteins increased and 21 proteins decreased by at least 3-fold in rmt2 cells relative to WT. The Cre number (Chlamydomonas gene accession number) for each protein and the fold change is shown. The fold change is given as Log2 values.

### Slide 12
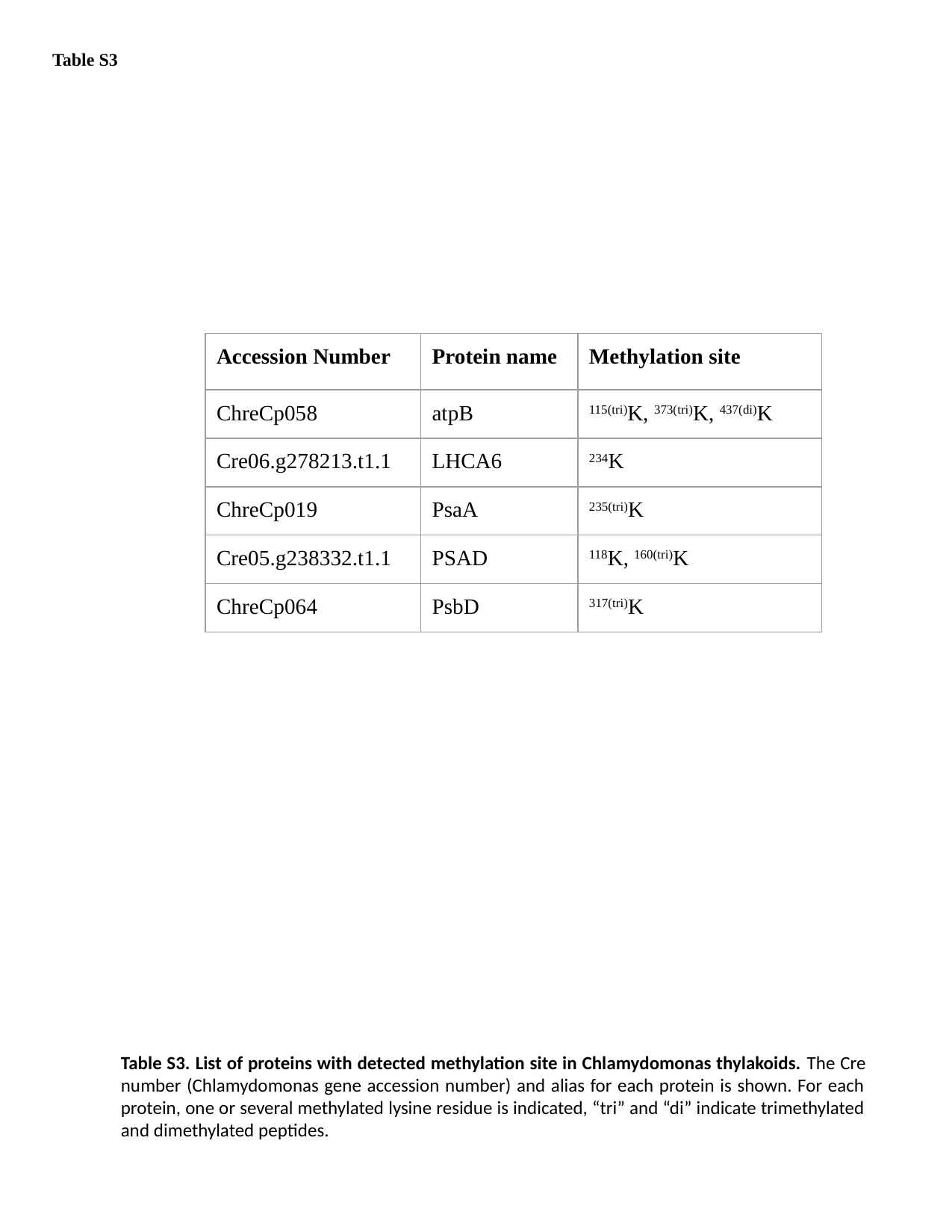

Table S3
| Accession Number | Protein name | Methylation site |
| --- | --- | --- |
| ChreCp058 | atpB | ^115(tri)^K, ^373(tri)^K, ^437(di)^K |
| Cre06.g278213.t1.1 | LHCA6 | ^234^K |
| ChreCp019 | PsaA | ^235(tri)^K |
| Cre05.g238332.t1.1 | PSAD | ^118^K, ^160(tri)^K |
| ChreCp064 | PsbD | ^317(tri)^K |
Table S3. List of proteins with detected methylation site in Chlamydomonas thylakoids. The Cre number (Chlamydomonas gene accession number) and alias for each protein is shown. For each protein, one or several methylated lysine residue is indicated, “tri” and “di” indicate trimethylated and dimethylated peptides.
